# Proteomics-constrained deconvolution reveals spatial cell-type programs in tumours

**DOI:** 10.64898/2026.06.01.729268

**Authors:** Esra Büşra Işık, Michael J. Haley, Ali Hussein Al-Anbaki, Leoma Bere, Federico Roncaroli, Karen Piper Hanley, Kevin Couper, David C Wedge, Robert Sellers, Alexander Baker, Pedro Oliveira, Jack Ashton, Robert G Bristow, Mauricio A. Álvarez, Sokratia Georgaka, Magnus Rattray

## Abstract

Accurately resolving cell-type mixtures in spatial transcriptomics remains challenging, particularly in heterogeneous tumours where cell populations are intermixed and matched single-cell references may be unavailable or poorly aligned. Current deconvolution approaches either require high-quality scRNA-seq references, suffer from scalability limitations, or lack interpretability. We introduce PISTACHIO, a proteomics-informed spatial transcriptomics deconvolution framework based on constrained non-negative matrix factorization with a negative-binomial likelihood. Rather than using probabilistic priors, PISTACHIO incorporates spatial cell-type constraints derived from paired Imaging Mass Cytometry, enforcing biologically grounded sparsity and explicit spatial feasibility of cell-type presence. PISTACHIO improved recovery of spatial cell-type distributions compared with Cell2location and STdeconvolve across synthetic and real tumour datasets. Our approach remains robust under cell-type assignment errors, maintaining high correlation with ground-truth under moderate noise, and achieves fast runtime on standard hardware, enabling practical large-scale deployment.

## 1 Introduction

Cellular behaviour is fundamentally shaped by spatial context. In multicellular tissues, cell identity, signalling activity and functional state emerge from interactions with neighbouring cells and the surrounding microenvironment. Spatially resolved molecular technologies have therefore become essential for studying tissue organisation in development, homeostasis and disease. Among these, spatial transcriptomics (ST) enables genome-wide measurement of RNA expression while preserving the spatial context of tissue sections, providing unprecedented insight into the spatial architecture of gene regulation and cellular heterogeneity within intact tissues (Ståhl et al. 2016; Liu et al. 2024; Bressan et al. 2023). However, most widely used ST platforms capture transcripts from spatial regions larger than individual cells, such that each spot typically contains RNA originating from multiple neighbouring cells. As a result, the observed gene expression profiles represent mixtures of multiple cell types, necessitating computational deconvolution to recover the underlying cellular composition.

A large class of deconvolution methods assumes that the expression profile at each spatial location can be represented as a linear combination of cell-type-specific transcriptional signatures derived from single-cell RNA sequencing (scRNA-seq) references. While these reference-based approaches have demonstrated strong performance when well-matched reference datasets are available, their accuracy can deteriorate when reference atlases are incomplete, noisy or derived from different biological conditions than the spatial samples under investigation (Sang-aram et al. 2024). In contrast, reference-free approaches attempt to infer latent expression programs directly from spatial data without relying on external references. Although these methods provide greater flexibility, the inferred factors are often difficult to interpret biologically and may not correspond directly to known cell identities (Saqib and Kim 2025). Consequently, accurately resolving the cellular composition of spatial transcriptomics measurements remains a central challenge in spatial omics analysis.

Another limitation of current spatial transcriptomics deconvolution frameworks is that they rely almost exclusively on RNA measurements, despite the increasing availability of spatial proteomics technologies capable of directly quantifying proteins in situ. Imaging-based proteomic platforms such as imaging mass cytometry (IMC) and multiplexed ion beam imaging enable simultaneous detection of dozens of protein markers at single-cell resolution within intact tissues (Giesen et al. 2014; Keren et al. 2018). Because proteins are the primary functional effectors of cellular processes and frequently serve as canonical markers for cell identity in histopathology and immunophenotyping, spatial proteomics provides information that is complementary to transcriptomic measurements. Importantly, mRNA abundance is often only moderately correlated with protein levels due to post-transcriptional regulation, variation in translation efficiency and differences in protein stability (Liu et al. 2016; Buccitelli and Selbach 2020). As a result, cell-type identification based solely on transcriptional measurements can be uncertain when marker genes exhibit weak, noisy or context-dependent expression patterns (Vogel and Marcotte 2012; Kleshchevnikov et al. 2022).

Beyond the limited integration proteomic information, many current deconvolution approaches primarily estimate cell-type proportions while treating gene-expression signatures as fixed reference profiles derived from single-cell RNA sequencing datasets (Andersson et al. 2020; Biancalani et al. 2021; Elosua-Bayes et al. 2021; Kleshchevnikov et al. 2022). However, the ability to recover cell-type-specific transcriptional programs directly from spatial data is critical for understanding how cellular states vary across tissue microenvironments and how tumour or immune programs evolve in situ. Methods such as DestVI and STdeconvolve have begun to address this challenge by modelling cell-state variation or latent transcriptional programs within spatial transcriptomic data (Lopez et al. 2022; Miller et al. 2022). Methods that can jointly infer both cell-type composition and transcriptional programs therefore offer an opportunity to connect spatial cell identity with functional gene-expression dynamics.

Recent advances in spatial multi-omics technologies have demonstrated the feasibility of jointly measuring RNA and protein expression within the same tissue sections. For example, spatial-CITE-seq enables simultaneous mapping of the entire transcriptome together with hundreds of protein markers at cellular resolution (Liu et al. 2023), while approaches such as DBiT-plus integrate spatial transcriptomics with multiplexed protein imaging to co-profile RNA and protein expression from the same tissue section (Liu et al. 2020). Despite these technological advances, most computational frameworks still analyse transcriptomic and proteomic measurements independently, and only limited efforts have attempted to incorporate spatial proteomic information as prior knowledge in spatial transcriptomics deconvolution (Ben-Chetrit et al. 2023; Koladiya et al. 2025; Chang et al. 2025). Consequently, spatial information on cell identity encoded in protein marker distributions is rarely used to constrain transcriptomic inference.

To address the above limitations, we introduce PISTACHIO, a proteomics-informed framework for spatial transcriptomics deconvolution that integrates spatial proteomic priors into matrix factorisation models (Fig. 1). PISTACHIO constrains non-negative matrix factorisation using binary spatial masks derived from paired IMC data, guiding each cell-type to locations where its protein markers are detected. The model employs a negative binomial likelihood to account for the overdispersed count distributions characteristic of spatial transcriptomics. By integrating proteomic priors with transcriptomic factorisation, PISTACHIO does not require matched scRNA-seq reference datasets, remains robust to moderate uncertainty in spatial masks, and simultaneously estimates both cell-type abundances and gene-expression programs. We benchmark PISTACHIO on synthetic datasets and apply it to glioblastoma and prostate cancer tissues with matched spatial transcriptomics and imaging mass cytometry measurements, demonstrating that proteomic constraints improve recovery of spatial cell distributions and enable biologically interpretable reconstruction of transcriptional programs.

**Fig. 1:**
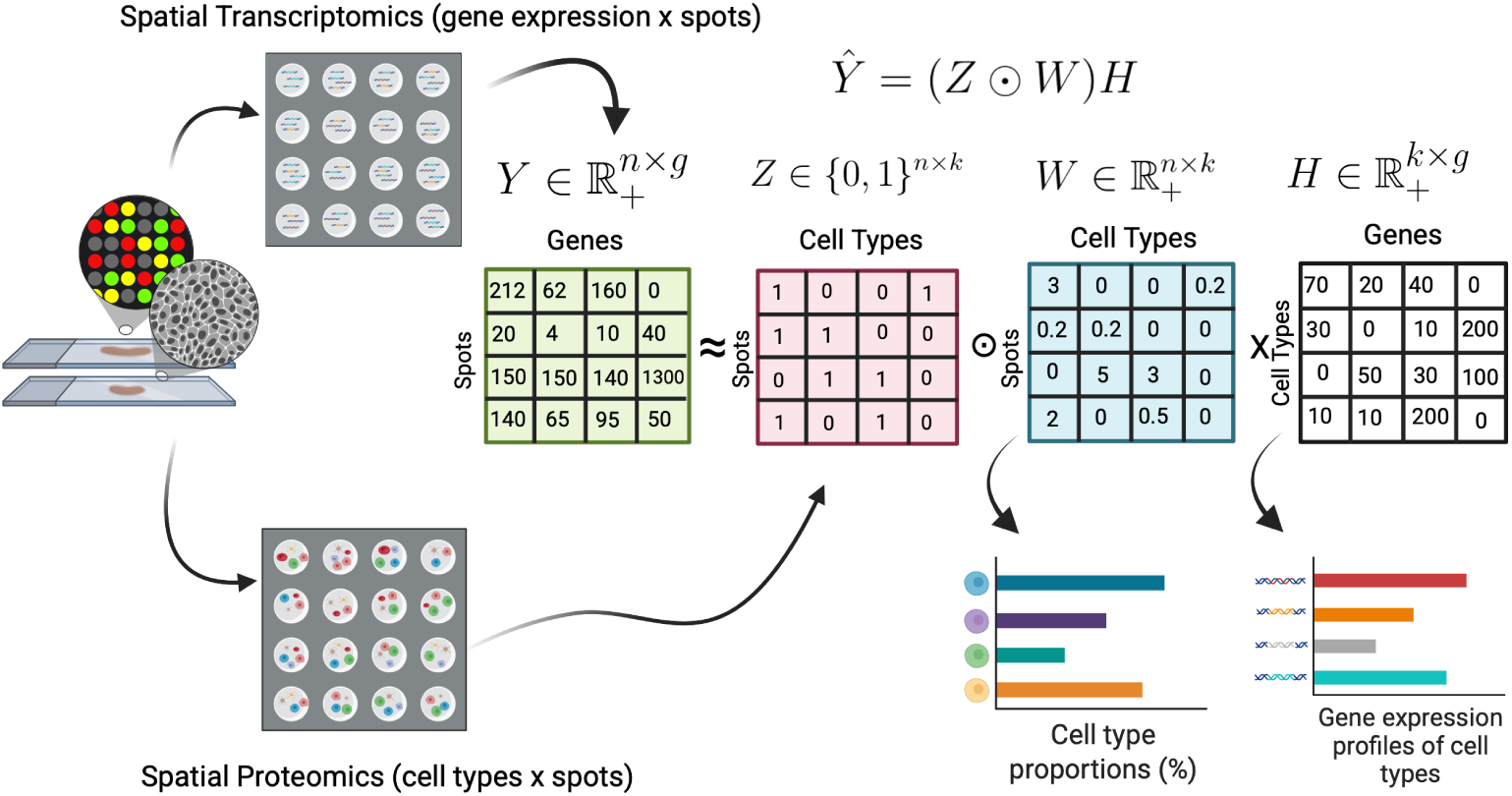
Overview of the PISTACHIO framework. Imaging mass cytometry is used to identify cell-types and generate a spatial cell-type presence mask. This mask constrains a non-negative matrix factorisation model applied to spatial transcriptomics counts, restricting each cell-type to locations where its protein markers are detected. The model decomposes the spatial transcriptomics matrix into a cell-type abundance matrix *W* and a gene-loading matrix *H*, representing spatial composition and cell-type transcriptional programs, respectively. A negative binomial objective is used to account for the overdispersed count structure of spatial transcriptomics data.

## 2 Results

### 2.1 Overview of PISTACHIO Framework

We formulate spatial transcriptomics deconvolution as a constrained Non-Negative Matrix Factorisation (NMF) problem guided by spatial proteomics priors. Let *Y* ∈ ℝ*^n^*^×*g*^ denote the observed gene expression counts matrix, where *n* is the number of spatial locations (spots) and *g* is the number of genes. We approximate *Y* as

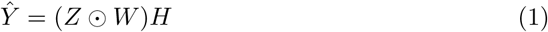

where 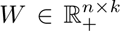 represents the abundance of *k* cell-types across spatial locations, 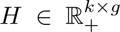 encodes their gene expression profiles, and *Z* ∈ {0, 1}*^n^*^×*k*^ is a binary spatial mask derived from paired IMC. The Hadamard product ⊙ enforces the spatial constraint such that *W_ik_* = 0 whenever *Z_ik_* = 0, thereby restricting each cell-type to biologically plausible spatial regions. Importantly, the mask encodes only the spatial feasibility of cell-type presence and not abundance.

To model the overdispersed nature of count-based transcriptomic data, PISTACHIO additionally employs a negative binomial NMF (NB-NMF) objective (Lyu et al. 2020), where *Ŷ*_*ij*_ is interpreted as the mean parameter *µ_ij_* of a negative binomial distribution. We parameterise the negative binomial distribution by its mean *µ_ij_* = *Ŷ*_*ij*_ and a gene-specific dispersion parameter *α_j_*. Up to additive constants, the negative log-likelihood is given by

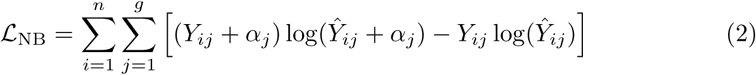

where *α* = (*α*_1_*, . . . , α_g_*) is a gene-wise dispersion vector broadcast across spatial locations. Following previous NB-NMF approaches (Lyu et al. 2020), we fixed *α_j_* = *α* = 10 for all genes, which provided stable performance across datasets. We also implement a Gaussian baseline to compare performance with the NB likelihood. Gaussian NMF was applied to normalised expression values, whereas NB-NMF was applied directly to raw count data. As a Gaussian baseline, we minimise the Frobenius reconstruction loss

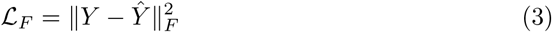

optimised using either coordinate descent or multiplicative update rules.

For NB-NMF, we use multiplicative update rules of the form

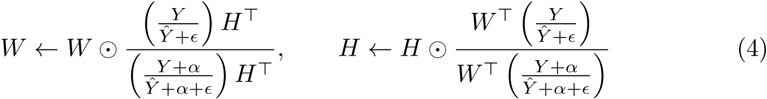

where *ɛ* is a small constant added for numerical stability to avoid division by zero and to ensure positivity of *Ŷ*. Analogous multiplicative updates are used for Gaussian NMF under the Frobenius loss, while coordinate descent is used as an alternative optimiser for the Gaussian objective.

The spatial constraint is enforced after initialisation and after each optimisation step by projecting

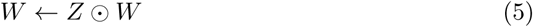

thereby guaranteeing that disallowed cell-types remain zero in forbidden spatial locations throughout training.

Optimisation is run for at most a fixed number of iterations and stopped early when the absolute change in the objective between successive iterations falls below a tolerance threshold. Convergence is monitored using the corresponding loss trajectory (Frobenius loss for Gaussian NMF and negative binomial loss for NB-NMF).

To resolve the scale ambiguity inherent to matrix factorisation, we apply a post-hoc normalisation after convergence:

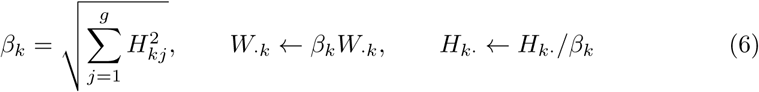

which rescales each component without changing the product *WH* and improves comparability across runs.

Initialisation schemes included random and scaled random initialisation proportional to gene-wise means. In all cases, the spatial mask was applied to the initial *W* to ensure feasibility, and all factor matrices were constrained to remain non-negative throughout optimisation. Reconstruction accuracy of the gene-expression matrix was assessed by comparing the observed counts *Y* with the reconstructed matrix *Ŷ* = *WH*. Performance was evaluated using complementary metrics, including root mean squared error (RMSE), mean absolute error (MAE), and Pearson correlation computed on the vectorised matrices. Pearson correlation was used as the primary measure of agreement.

### 2.2 Proteomics-constrained deconvolution accurately recovers spatial composition and transcriptional programs in synthetic data

To evaluate the performance of PISTACHIO in a controlled setting, we first tested the method on synthetic spatial transcriptomics data where the underlying ground-truth is known. The dataset was generated from single-donor human glioblastoma (GBM) scRNA-seq profiles, allowing both spot-level cell-type contributions and cell-type-specific gene expression programs to be measured directly (see Methods). Using its negative binomial formulation, PISTACHIO reconstructed the observed spatial transcriptomics data with high fidelity (Fig. 2a). This indicates that the factorisation captures the main structure of the gene expression signal across spatial locations.

**Fig. 2:**
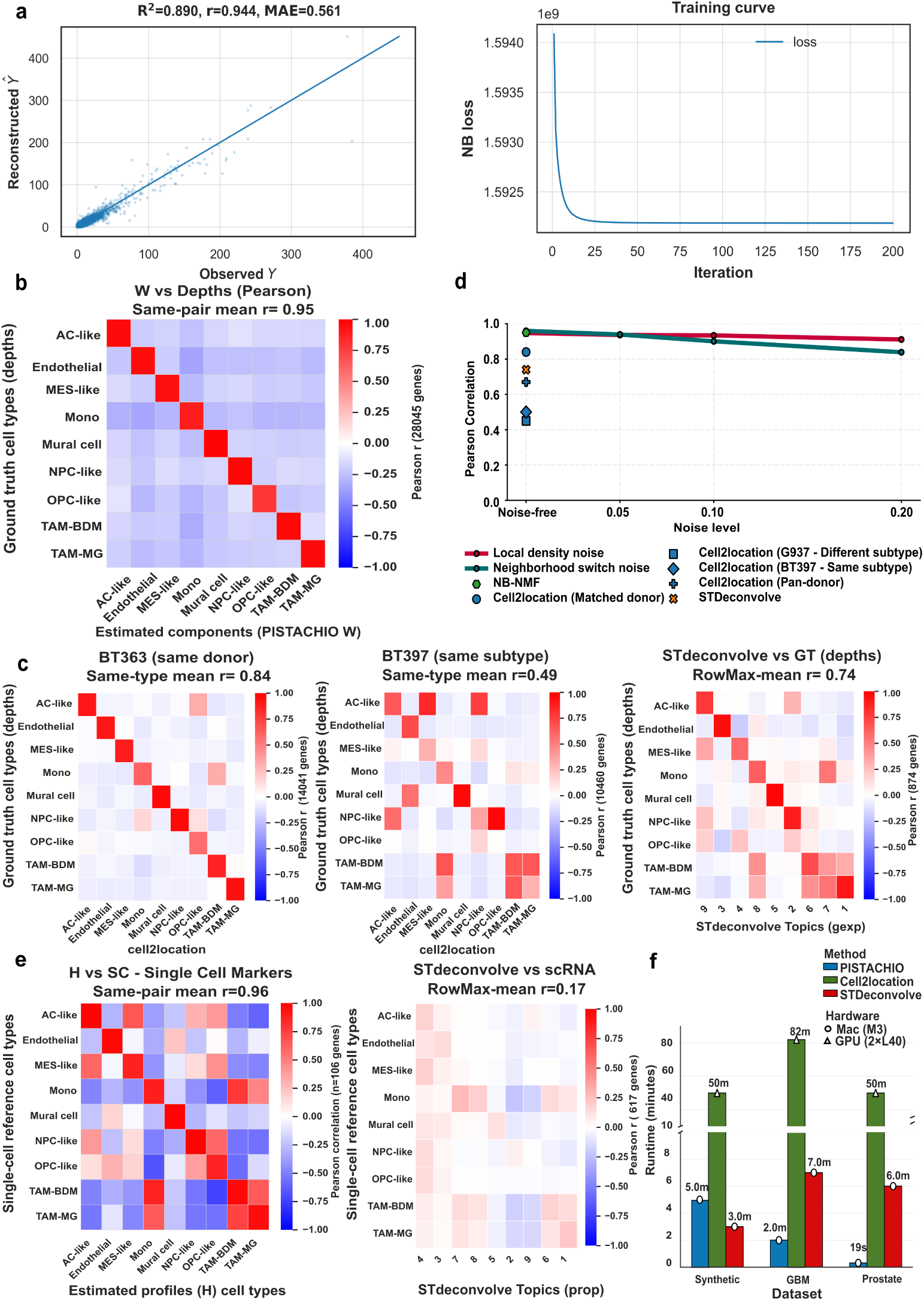
Benchmarking of PISTACHIO on synthetic spatial transcriptomics data. **(a)** Reconstruction of observed spatial transcriptomics counts using the NB-NMF model. Left, observed versus reconstructed counts. Right, negative binomial training loss across optimisation iterations. **(b)** Pearson correlation between inferred cell-type abundances *W* and ground-truth depth contributions *D* (UMI-based cell-type contributions per spot) across spatial locations. **(c)** Comparison of abundance recovery across methods. Pearson correlation heatmaps between inferred abundances and ground-truth for Cell2location (donor-matched BT363 reference and BT397 reference) and STdeconvolve. **(d)** Robustness to noise in the spatial prior. Same-type Pearson correlation between inferred and ground-truth abundances across increasing perturbation levels for two spatial noise models. **(e)** Recovery of gene-expression programs. Pearson correlation between inferred gene-loading profiles *H* and single-cell reference signatures using marker genes, compared with STdeconvolve topic programs. **(f)** Runtime comparison across datasets (synthetic, GBM and prostate) for PISTACHIO, Cell2location and STdeconvolve.

More importantly, the inferred spatial abundance matrix *W* closely matched the true cell-type contributions across spots. Here, ground-truth cell-type contributions are defined as the UMI-based depth matrix *D*, representing the total transcript counts contributed by each cell-type within each spot. The correlation matrix between inferred and ground-truth abundances showed a clear diagonal pattern with very little off-diagonal signal (same-pair mean *r* ≈ 0.95; Fig. 2b), indicating that individual cell-types were correctly localised across the tissue without substantial cross-type mixing. This behaviour was consistent across different ways of handling zero-valued entries in the correlation analysis: excluding or retaining shared zero entries yielded similarly strong agreement with the ground-truth (Supplementary Fig. S1a). Scatter plots for individual cell-types further confirmed a near-linear relationship between inferred and true abundances across a wide range of values (Supplementary Fig. S1a).

We next compared PISTACHIO with two representative deconvolution approaches. Cell2location uses scRNA-seq reference profiles to guide inference, whereas STdeconvolve attempts to identify latent cell-type programs directly from spatial data without prior information. When provided with a well-matched scRNA-seq reference, Cell2location recovered cell-type abundances with reasonably high accuracy (same-type mean *r* ≈ 0.84; Fig. 2c). However, this performance declined substantially when the reference profiles were replaced with data from different donors or pooled datasets (Supplementary Fig. S1d), highlighting the sensitivity of reference-based approaches to inter-patient transcriptional variation. In contrast, STdeconvolve identified latent topics that only partially corresponded to individual cell-types, with several topics representing mixtures of multiple populations and no clear one-to-one correspondence with the ground-truth composition (Fig. 2c).

Because PISTACHIO relies on spatial priors derived from proteomics measurements, we next asked how sensitive the model is to inaccuracies in these priors. To test this, we introduced structured perturbations to the binary cell-type presence mask that simulates realistic sources of error such as reduced detection sensitivity or spatial mixing at cell boundaries. Across both noise models, PISTACHIO remained highly stable (Fig. 2d). At moderate noise levels (5–10%), the agreement between inferred and true cell-type abundances decreased only slightly, and even under stronger perturbations (20% noise) the model retained strong correspondence with the ground-truth. These results suggest that spatial constraints guide the model without rigidly determining the solution, allowing the method to tolerate substantial uncertainty in proteomics-derived priors.

Beyond recovering spatial cell-type composition, PISTACHIO also infers cell-type-specific transcriptional programs through the gene-loading matrix *H*. We therefore examined whether the inferred programs correspond to known cell-type expression signatures. Comparing the inferred profiles with single-cell reference signatures revealed very strong agreement when focusing on informative marker genes (same-pair mean *r* ≈ 0.96; Fig. 2e). The correlation matrix displayed a clear one-to-one alignment between inferred and reference profiles, indicating that the model captures transcriptional identities rather than simply fitting the observed counts. Off-diagonal similarities were largely restricted to closely related lineages, such as TAM-BDM and TAM-MG populations, which are known to share partially overlapping transcriptional states in glioblastoma.

In contrast, gene programs inferred by STdeconvolve showed much weaker correspondence with single-cell reference profiles (row-max mean *r* ≈ 0.17; Fig. 2e), reflecting the difficulty of interpreting unsupervised topic models in terms of specific cell identities. Importantly, the strong agreement observed for PISTACHIO was not limited to externally defined marker genes. Marker sets derived directly from the inferred *H* profiles produced similarly high correspondence with single-cell reference signatures (Supplementary Fig. S1a), indicating that the recovered transcriptional programs are internally consistent and biologically meaningful.

Finally, PISTACHIO was also substantially faster in practice, completing inference roughly an order of magnitude faster than Cell2location across the synthetic benchmarks, as well as the real datasets introduced in the subsequent sections (Fig. 2f).

### 2.3 Robustness analyses confirm stable deconvolution under imperfect priors and model assumptions

We next examined whether the performance of PISTACHIO is strongly dependent on the accuracy of the spatial prior or on the specific statistical assumptions of the model. To address this, we performed two complementary analyses that probe the robustness of the method.

First, we tested whether PISTACHIO can resolve the hidden structure when prior information is deliberately simplified. In this “unknown subtype” experiment, pairs of biologically related cell-types were merged into a single “Unknown” category in the spatial prior while keeping the full spot-level expression counts unchanged (see Methods). Then, two components were allocated to this collapsed category such that the model had to separate the underlying subtypes using transcriptional information alone.

Across all tested scenarios, including highly similar immune subtypes as well as more distinct vascular and malignant populations, PISTACHIO successfully separated the hidden cell-types and recovered a clear one-to-one correspondence with the ground-truth (same-pair Fisher-averaged *r* ≈ 0.95–0.96; Supplementary Fig. S1c). These results indicate that the method performs genuine deconvolution rather than simply assigning cell-types based on the spatial prior, and can exploit transcriptional differences to resolve latent substructure even when prior annotations are incomplete.

We then examined the importance of explicitly modelling count overdispersion in spatial transcriptomics data. To this end, we compared the negative binomial NMF formulation used in PISTACHIO with Gaussian NMF baselines optimised using coordinate descent and multiplicative updates. Although the Gaussian models were able to capture some aspects of the data structure, their performance was consistently weaker and less stable than the negative binomial formulation (Supplementary Fig. S2a,b). In particular, Gaussian NMF showed reduced agreement with the ground-truth cell-type abundances (same-pair Fisher-averaged *r* ≈ 0.6–0.95, depending on the optimiser) and substantially poorer recovery of gene-expression programs when applied to raw count data with all the genes. Multiplicative updates improved performance relative to coordinate descent by achieving lower reconstruction loss under Gaussian assumptions (Supplementary Fig. S2c) and after filtering the genes based on differentially expressed ones, but still failed to match the accuracy of the negative binomial model.

### 2.4 Accurate recovery of cell-type composition and spatial organization in prostate tissue

We next evaluated the performance of PISTACHIO on two localised prostate cancer tissue sections (ST2 and ST1) annotated for histologic elements of normal stroma or prostate epithelium versus malignant adenocarcinoma or the sub-histology intraductal carcinoma/cribiform architecture (IDC/CA). Spatial transcriptomics has been used to infer differential gene expression between normal and malignant prostate epithelium, but has not been aligned with parallel spatial proteomics (Erickson et al. 2022; Ali et al. 2026). We therefore utilised matched Visium spatial transcriptomics and Imaging Mass Cytometry (IMC) to annotate the two specimens. IMC-derived cell annotations revealed heterogeneous and sparse cell-type distributions across spatial spots (Fig. 3a). Several epithelial and stromal populations, including epithelial tumour cells and CAF/myofibroblasts, were broadly distributed across both samples, whereas immune populations such as NK-like cytotoxic cells and myeloid APC populations appeared only in subsets of spatial locations, reflecting the heterogeneous cellular composition of the tumour microenvironment.

**Fig. 3:**
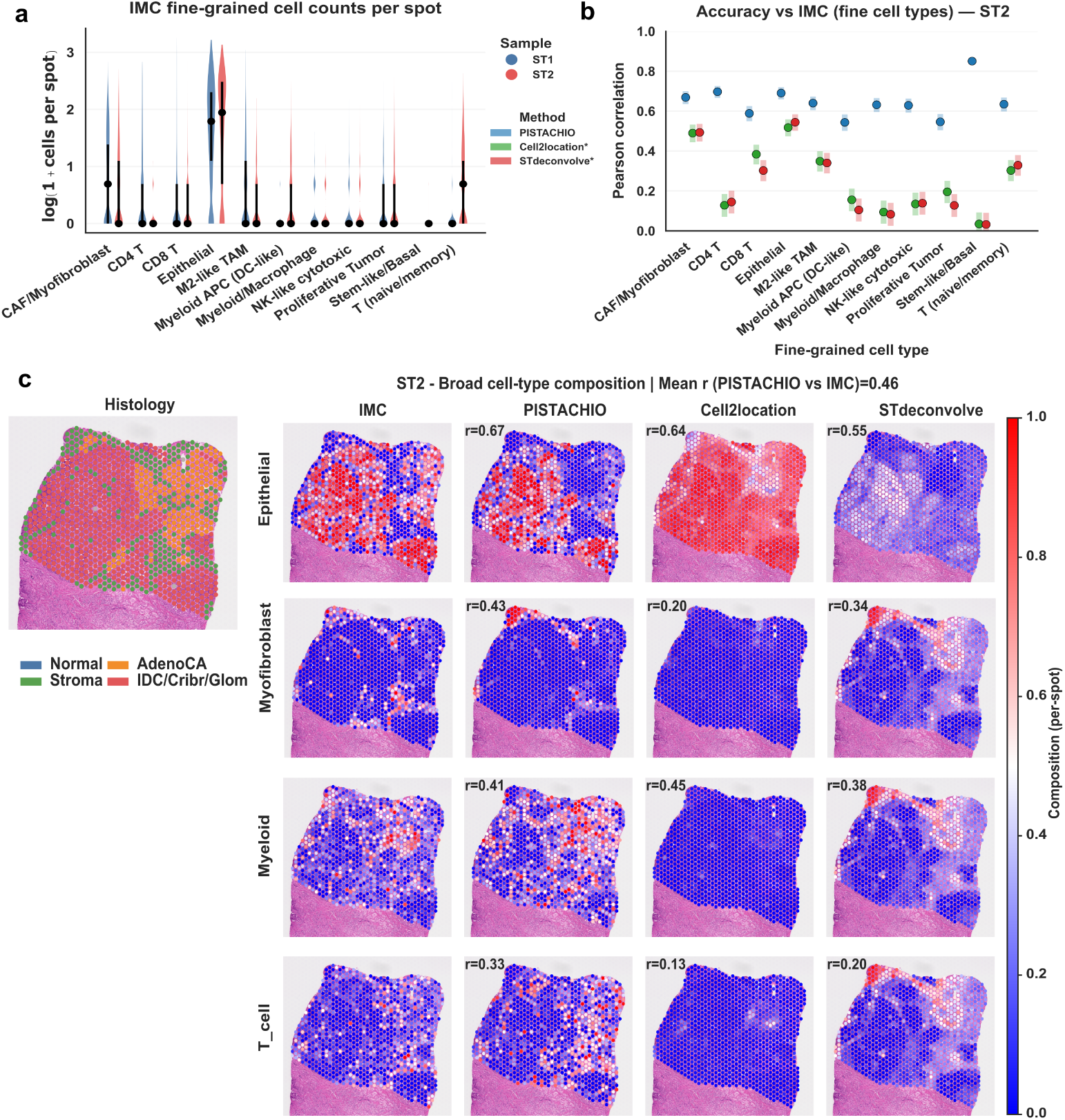
Recovery of spatial cell-type composition in prostate tissue. **(a)** Distribution of IMC-derived cell counts per Visium spot across fine-grained cell-types for prostate sections ST1 and ST2. **(b)** Pearson correlation between inferred cell-type abundances and IMC-derived area-based ground-truth for fine-grained cell-types in sample ST2, comparing PISTACHIO, Cell2location and STdeconvolve. **(c)** Spatial comparison of broad cell-type composition in sample ST2. Columns show histological annotation, IMC-derived ground-truth and spatial maps inferred by PISTACHIO, Cell2location and STdeconvolve. Rows correspond to epithelial, myofibroblast, myeloid and T cell compartments. Colour scales indicate per-spot cell-type proportions.

We first evaluated the recovery of fine-grained cell-type composition by comparing inferred cell-type abundances *W* with IMC-derived area-based ground-truth, defined as the summed cellular area occupied by each cell-type within each Visium spot. In sample ST2, PISTACHIO showed higher agreement with IMC measurements than Cell2location and STdeconvolve across the majority of cell-types (Fig. 3b). While Cell2location recovered several epithelial and stromal populations, correlations were reduced for multiple cell-types, and STdeconvolve topics showed weaker correspondence to biologically defined cell identities.

Inspection of the full correlation matrices further illustrates these differences. For both ST2 and ST1, correlations between PISTACHIO-inferred abundances and IMC-derived cell-type densities displayed a pronounced diagonal structure, indicating correspondence between inferred and reference cell-types (Supplementary Fig. 3a,b). In contrast, correlation matrices for Cell2location and STdeconvolve showed weaker diagonal enrichment and increased off-diagonal signal, consistent with greater cross-type mixing in the inferred compositions (Supplementary Fig. 3c).

Because many fine-grained cell-types are absent from a large fraction of spatial locations, we additionally examined the effect of including or removing known zero entries (available from ground-truth IMC-derived cell-types) on correlation estimates. Including zero entries within correlation coefficient calculations increased correlations for all methods, whereas removing them produced more conservative estimates that emphasise agreement within spatially informative regions (Supplementary Fig. 3a,b).

We next examined whether the inferred abundances preserved tissue-level spatial organisation. In sample ST2, spatial maps inferred by PISTACHIO recapitulated epithelial, myofibroblast, myeloid and T cell compartments observed in IMC measurements (Fig. 3c). Similar spatial patterns were observed in the independent section ST1 (Supplementary Fig. 4). In comparison, Cell2location produced more diffuse spatial distributions, while STdeconvolve topics showed weaker correspondence to histologically defined structures. Notably, STdeconvolve topic 3 correlated with multiple biologically distinct populations (myofibroblasts, myeloid cells and T cells), indicating that a single latent topic captured mixed biological signals rather than resolving individual cell-types.

### 2.5 Improved estimation of cell-type-specific expression profiles with interpretable cell-type contributions

Beyond recovering spatial cell-type composition, PISTACHIO estimates cell-type-specific transcriptional programs (*H*) and enables decomposition of spatial gene expression into interpretable cell-type contributions. We evaluated the biological validity of these inferred programs using proteomic markers, external scRNA-seq references, and spatial reconstruction of canonical tumour and stromal genes.

We first examined IMC protein marker profiles associated with the fine-grained cell-type annotations used to construct spatial constraints in the model (Fig. 4a; Supplementary Fig. 5a). These marker profiles delineate epithelial, stromal and immune compartments of the tumour microenvironment and provide an independent reference for interpreting the inferred transcriptional programs.

**Fig. 4:**
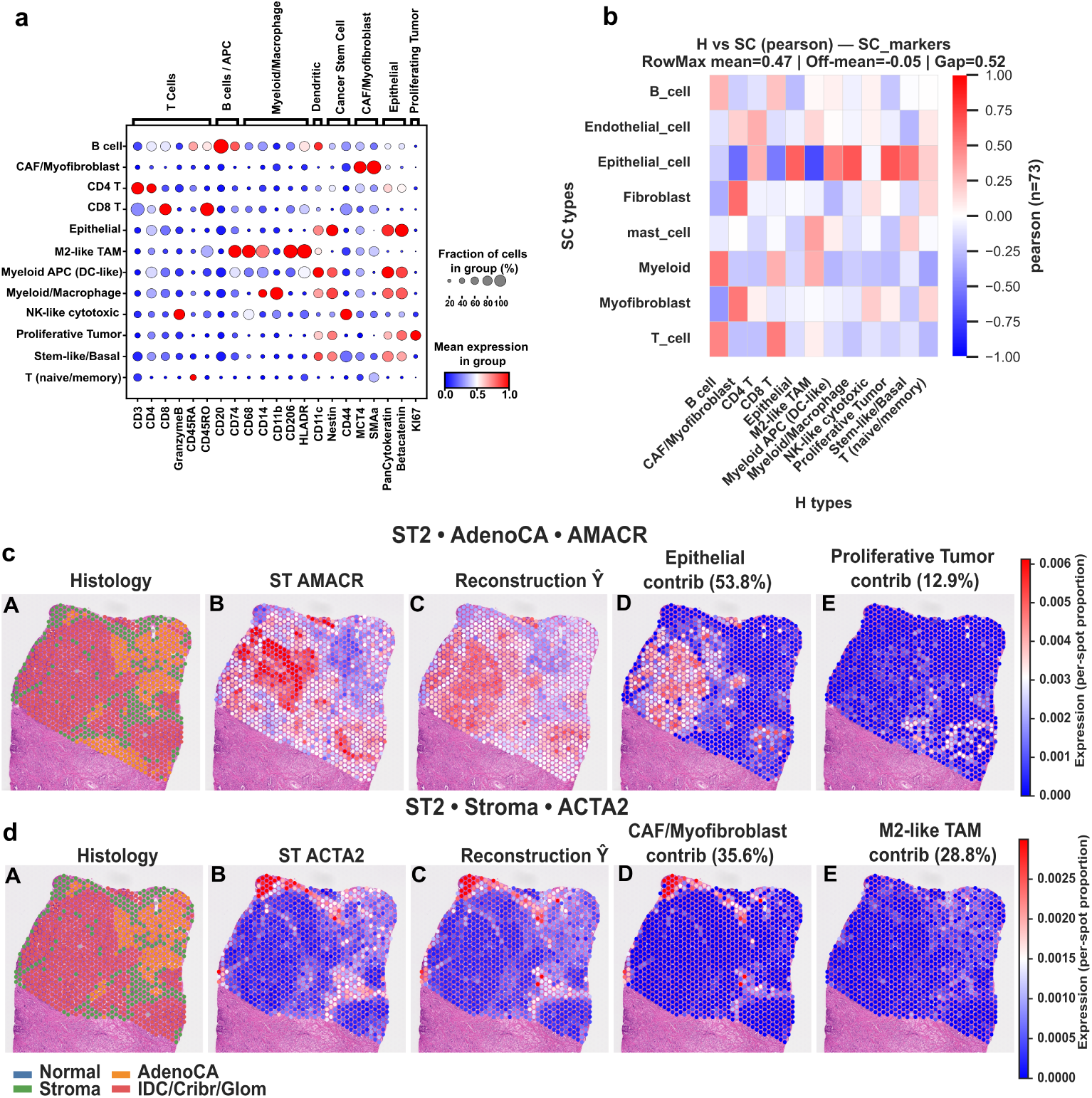
Reconstruction of cell-type transcriptional programs and gene-level spatial contributions. **(a)** IMC protein marker profiles defining fine-grained cell-types used to guide spatial constraints. Dot size indicates the fraction of cells expressing each marker within a cell-type and colour indicates mean protein expression. **(b)** Correspondence between inferred transcriptional programs (*H*) and external scRNA-seq reference cell-types using curated marker genes. Heatmap shows Pearson correlation between inferred programs and reference cell-type profiles. **(c)** Spatial reconstruction of the tumour-associated marker gene *AMACR* in prostate section ST2. Columns show histology annotation, observed spatial transcriptomics signal, reconstructed expression *Ŷ*, and dominant cell-type contributions. **(d)** Spatial reconstruction of the stromal marker *ACTA2* in the same section, shown as in (c). Colour scales indicate per-spot expression after log transformation and percentile clipping.

We next compared the inferred transcriptional programs (*H*) with cell-type expression profiles from an external prostate scRNA-seq reference using curated marker genes (Fig. 4b; Supplementary Fig. 5b). In both samples, inferred programs showed correspondence with known cellular lineages. In sample ST2, the mean row-wise maximum correlation between inferred programs and single-cell references was approximately 0.44 with low off-diagonal similarity (off-mean ∼ −0.04). A comparable pattern was observed in sample ST1, where the mean row-wise maximum correlation reached 0.45. Agreement was strongest for cell-types with well-defined proteomic markers, such as epithelial and stromal fibroblast populations, whereas immune and myeloid populations showed broader similarity across related lineages. This likely reflects both biological similarity among immune states and the challenges of resolving mixed transcriptomic signals within spatial spots (Liu et al. 2016). In addition, the available single-cell reference does not match perfectly the spatial samples analyzed here, which may further limit the achievable correspondence.

We next examined whether PISTACHIO provides interpretable spatial attribution of gene expression by analysing two well-established prostate biomarker genes: the prostate tumour marker *AMACR* and the stromal activation marker *ACTA2* (Rubin et al. 2002; Nurmik et al. 2020). For both genes, reconstructed spatial expression (*Ŷ*) closely reproduced the observed spatial transcriptomics signal (Fig. 4c,d; Supplementary Fig. 5c).

In sample ST2, *AMACR* expression was enriched within tumour-associated epithelial regions and was primarily attributed to epithelial cells (53.8% of reconstructed signal), with an additional contribution from proliferative tumour cells (12.9%) (Fig. 4c). In contrast, *ACTA2* expression was concentrated in stromal compartments and was predominantly explained by CAF/myofibroblast populations (35.6%), with a substantial contribution from M2-like tumour-associated macrophages (28.8%) (Fig. 4d).

Comparable patterns were observed in the independent section ST1 (Supplementary Fig. 5). In this sample, *AMACR* expression was again primarily attributed to epithelial cells (44.9%), whereas *ACTA2* expression was dominated by CAF/myofi-broblast populations (46.1%) with a smaller epithelial contribution (13.4%).

To further contextualise these results, we compared transcriptional programs inferred by PISTACHIO with those obtained from STdeconvolve, which also learns gene expression topics directly from spatial transcriptomics data. STdeconvolve topics showed mixed correspondence with scRNA-seq profiles and weaker agreement with IMC-defined cell identities (Supplementary Fig. 3c), whereas PISTACHIO programs showed clearer alignment with both proteomic markers and spatial cell-type annotations. Cell2location could not be evaluated in this analysis because it does not infer cell-type-specific gene expression programs, instead relying on predefined reference signatures. Together, these results indicate that integrating proteomic spatial constraints improves the biological specificity of inferred transcriptional programs while preserving interpretability at the gene level.

### 2.6 Program-level differences reveal cell-type-specific transcriptional changes between samples

To characterise transcriptional differences between the two prostate sections (ST2 and ST1) in a cell-type-resolved manner, we summarised the inferred gene-loading matrix *H* into a panel of biologically interpretable programs representing tumour identity, lineage state, metabolic stress, immune activity and signalling (Supplementary Fig. 6). Program scores were computed from mean gene-wise z-scored *H* loadings within each cell type, and cross-sample shifts were quantified as 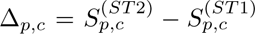 gene-level sign-flipping bootstrap with Benjamini–Hochberg correction (Methods).

Across the inferred programs, transcriptional differences between sections were strongly cell-type specific rather than global (Fig. 5a; Supplementary Fig. 7). Similarly, Proliferative Tumor cells displayed high scores for the Proliferation / Cell Cycle program across samples, reflecting expression of canonical cell-cycle genes such as *MKI67* and *TOP2A* that mark actively cycling tumour populations (Song et al. 2022). These observations indicate that core malignant transcriptional states are largely preserved across sections.

**Fig. 5:**
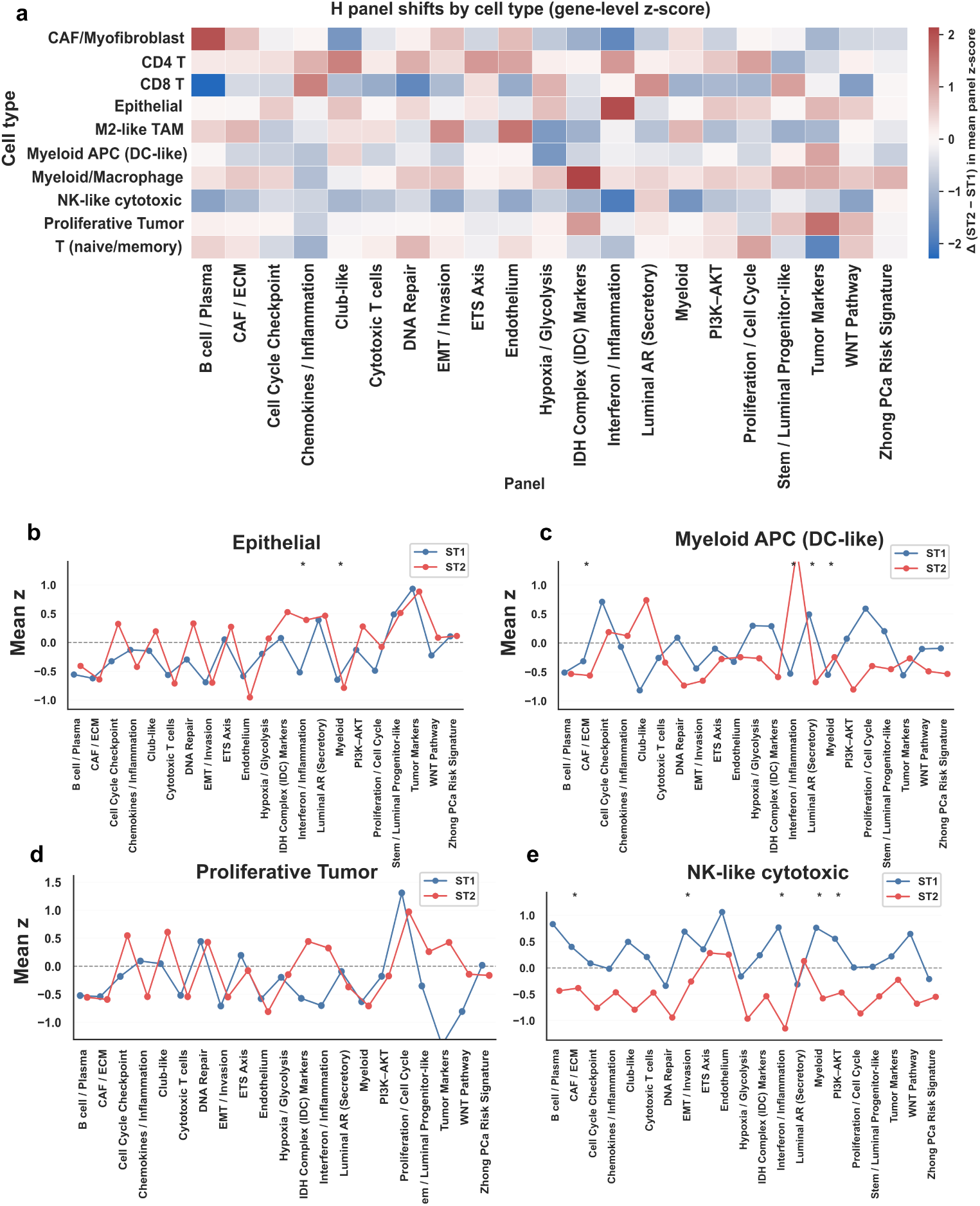
Program-level transcriptional differences between prostate tumour samples. **(a)** Heatmap showing differences in inferred transcriptional program activity between prostate sections ST2 and ST1 across cell-types and gene programs. Values represent the change in mean program z-score between samples (Δ, ST2–ST1). **(b–e)** Program activity profiles for selected cell-types comparing samples ST1 (blue) and ST2 (red). Each point represents the mean z-scored activity of a gene program within the indicated cell type. Panels show epithelial tumour cells (**b**), myeloid APC (DC-like) cells (**c**), proliferative tumor cells (**d**), and NK-like cytotoxic cells (**e**). Asterisks indicate statistically significant differences based on bootstrap testing.

Immune populations displayed more heterogeneous transcriptional changes. Myeloid APC (DC-like) cells showed increased Interferon / Inflammation activity in ST2 together with enrichment of myeloid-associated programs (Fig. 5b,c), consistent with an activated antigen-presenting phenotype within inflamed tumour microenvironments (Owen et al. 2020; Kwon et al. 2021). In contrast, NK-like cytotoxic cells exhibited reduced inflammatory and myeloid-associated signatures, indicating distinct activation states among cytotoxic lymphocyte populations.

T cell populations showed selective shifts between sections. CD4 T and T (naive/memory) cells in ST2 displayed increased Proliferation / Cell Cycle activity, whereas CD8 T cells showed comparatively limited transcriptional differences (Supplementary Fig. 7, Supplementary Table 1), suggesting that proliferative expansion occurs preferentially within specific T cell subsets.

Stromal compartments also exhibited transcriptional remodelling. CAF/Myofibroblast populations in ST2 showed enrichment of extracellular matrix and stromal programs together with reduced interferon-associated activity, consistent with the role of cancer-associated fibroblasts in regulating extracellular matrix organisation and stromal signalling within prostate tumours (Kalluri 2016). Among macrophage-lineage populations, M2-like TAM cells showed reduced inflammatory and proliferative programs, whereas broader Myeloid/Macrophage populations exhibited more modest changes, reflecting the functional plasticity of tumour-associated macrophages (Mantovani et al. 2017).

These analyses demonstrate that PISTACHIO-derived transcriptional programs capture biologically meaningful and cell-type-specific variation between prostate sections. While core malignant transcriptional identities remain largely stable, differences between samples are primarily driven by immune activation and stromal remodelling programs across the entire tumour microenvironment.

### 2.7 Proteomics-constrained deconvolution recovers broad spatial organisation across glioblastoma samples

We next evaluated the ability of PISTACHIO to recover spatial cell-type composition in glioblastoma (GBM), a tumour characterised by marked intra- and inter-tumoural heterogeneity, diffuse infiltration of adjacent brain, and complex, overlapping tumour, immune and stromal cell phenotypes and functional cell states. Across the four GBM tissue sections (A–D), inferred broad cell-type composition varied substantially between cases, with differences in the relative abundance of astrocyte-like, mesenchymal-like, NPC-like and OPC-like populations, together with myeloid immune and vascular/stromal populations (Fig. 6a). These patterns indicate that the model captured sample-specific ecosystem structure rather than converging to a common composition profile.

**Fig. 6:**
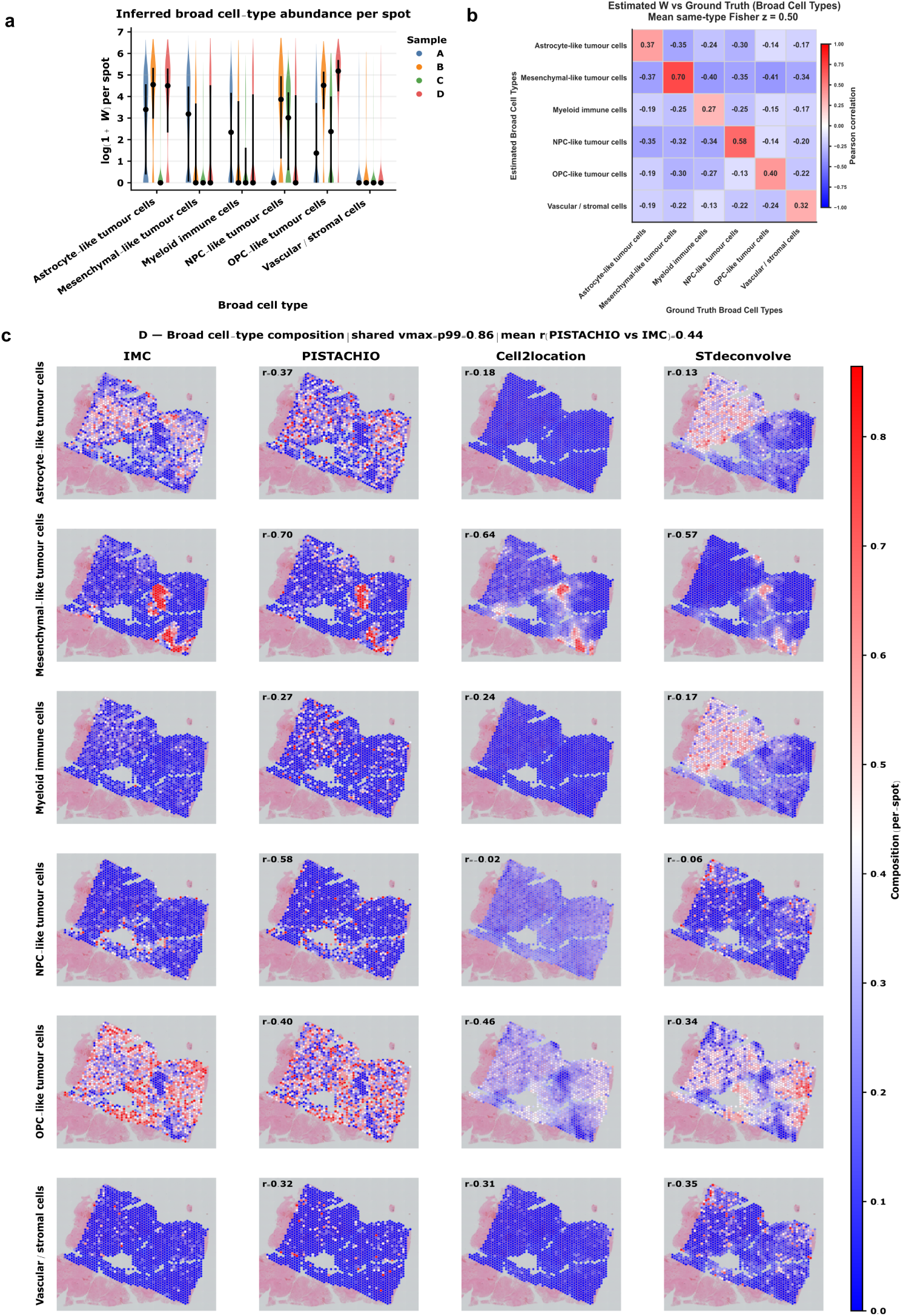
Recovery of spatial cell-type composition in glioblastoma. **(a)** Distribution of inferred broad cell-type abundance per spot across GBM samples A–D. **(b)** Pearson correlation between inferred broad cell-type abundances and IMC-derived broad ground-truth in sample D. **(c)** Spatial comparison of broad cell-type composition in sample D. Columns show IMC-derived ground-truth, spatial maps inferred by PISTACHIO, Cell2location and STdeconvolve. Rows correspond to astrocyte-like tumour cells, mesenchymal-like tumour cells, myeloid immune cells, NPC-like tumour cells, OPC-like tumour cells and vascular/stromal cells. Colour scales indicate per-spotbroad cell-type composition.

We first compared inferred fine-grained cell-type abundances with IMC-derived ground-truth. Across samples A–C, correlation matrices between inferred *W* and IMC-derived spot-level cell-type depths showed clear diagonal enrichment, indicating recovery of many fine-grained populations with the expected one-to-one correspondence (Supplementary Fig. 9a–c). Agreement was strongest for tumour cells and several stromal or vascular populations with relatively distinctive spatial patterns, whereas closely related immune and transitional states showed weaker separation. Including shared zero–zero entries increased same-pair correlations, while excluding shared zeros provided a more stringent evaluation focused on spatially informative regions; under both settings, diagonal structure remained evident (Supplementary Fig. 8a,b). IMC protein marker profiles were also consistent with the biological coherence of the fine-grained cell-type definitions used to construct the masks (Supplementary Fig. 8c).

Because fine-grained evaluation is particularly challenging in GBM, we next examined recovery at the level of broad biological compartments. In sample D, correlations between inferred broad abundances and IMC-derived broad ground-truth showed a clear diagonal pattern with limited off-diagonal correspondence (Fig. 6b). Agreement was strongest for mesenchymal-like (*r* = 0.70) and NPC-like (*r* = 0.58), followed by OPC-like (*r* = 0.40), and astrocyte-like cell phenotypes and functional states (*r* = 0.37), and vascular/stromal cells (*r* = 0.32). The spectrum of myeloid cells was recovered more modestly (*r* = 0.27), consistent with the spatial diffuseness and phenotypic overlap of these populations in GBM. Notably, most off-diagonal correlations were negative, indicating that inferred components retained lineage specificity rather than capturing non-specific spatial signal.

We then compared PISTACHIO with Cell2location and STdeconvolve using the same broad compartment framework. In sample D, PISTACHIO reproduced the major spatial distributions seen in IMC-derived maps, including focal mesenchymal-like regions, spatially restricted NPC-like cells, and broader OPC-like and vascular/stro-mal patterns (Fig. 6c). Cell2location recovered some large-scale trends, particularly for mesenchymal-like cells, but generally produced smoother maps with reduced correspondence for astrocyte-like, myeloid and NPC-like compartments. STdeconvolve captured part of the global spatial variation but showed weaker agreement with biologically defined compartments. Quantitatively, PISTACHIO achieved higher mean same-type broad-compartment correlation in sample D (mean Fisher *z* = 0.50) than Cell2location (mean Fisher *z* = 0.15) or STdeconvolve (mean Fisher *z* = 0.16) (Fig. 6b,c).

The same overall behaviour was observed across the remaining GBM samples. In samples A–C, PISTACHIO recovered the dominant broad spatial architectures visible in IMC-derived ground-truth, with consistent diagonal enrichment in broad-compartment correlation matrices and spatial maps that were closer to the IMC-derived reference (Supplementary Fig. 9a–c; Supplementary Fig. 10a–c). Performance varied between sections, with sample C showing the strongest broad-compartment agreement and samples A and B showing more moderate recovery, particularly for the astrocyte-like and vascular/stromal compartments. These results indicate that proteomics-constrained deconvolution preserves the major spatial ecosystem structure of GBM despite is considerable heterogeneity.

### 2.8 Functional annotation reveals biologically coherent GBM transcriptional programs

We next examined whether the inferred gene-loading matrix *H* captured biologically meaningful transcriptional programs in glioblastoma. In contrast to the synthetic benchmark and the prostate carcinoma cohort, direct correspondence between inferred GBM transcriptional programs and external scRNA-seq reference profiles was only moderate when evaluated across the full set of fine-grained IMC-derived cell-types (Supplementary Fig. 11a). The correlation matrix showed partial lineage structure, but lacked the strong one-to-one alignment observed in the synthetic data, with overlap among malignant, immune and transitional states.

To determine whether this weaker correspondence partly reflected imperfect cross-modal annotation rather than failure of the inferred programs themselves, we used MaxFuse to establish higher-confidence correspondences between protein-defined and RNA-defined cell states. The resulting protein–RNA matching matrix showed consistent mapping of several IMC-defined populations onto canonical RNA cell classes, including astrocyte-like, mesenchymal-like, NPC-like, OPC-like, endothelial, mural and TAM-associated phenotypes and functional states, whereas hybrid and transitional populations mapped across related RNA lineages (Supplementary Fig. 12). Re-evaluating *H* using these matched RNA cell classes improved the apparent single-cell correspondence (Supplementary Fig. 11b), with clearer agreement for MES-like, NPC-like, OPC-like, radial glia-like and TAM-related programs. These results indicate that the inferred GBM transcriptional programs retain lineage information, but that their interpretation is limited by the incomplete equivalence between protein-based and RNA-based annotation systems in tumours with high cell plasticy.

We therefore next evaluated the inferred *H* matrix using functional annotation. Broad cell-type composition differed substantially across the four GBM samples (Supplementary Fig. 13a), suggesting that each section was dominated by a distinct combination of tumour ecosystems and microenvironmental states. Consistent with this, Hallmark enrichment analysis revealed recurrent signatures strongly aligned with established GBM biology (Supplementary Fig. 13b). Enriched terms included hypoxia, TNF-alpha signaling via NF-kB, interferon alpha response and interferon gamma response, among other related terms.

The dominant enrichments also differed between samples. Sample A showed prominent hypoxia-associated and inflammatory hallmarks, consistent with a programme shaped by metabolic stress and immune activation. Sample B showed stronger enrichment for oxidative phosphorylation, reactive oxygen species, interferon-related signalling and TGF-*β*-associated pathways, suggesting a distinct balance between metabolic and inflammatory programmes. Sample C displayed enrichment of hypoxia, TNF/NF-kB signalling and epithelial–mesenchymal transition, whereas sample D showed stronger enrichment of angiogenesis, coagulation, myogenesis and mesenchymal-remodelling, consistent with a more vascularised or stromally remodelled microenvironment.

Gene Ontology enrichment supported the biological relevance of the inferred transcriptional programs. Enriched biological processes reflected both neural lineage features (for example neurogenesis and synaptic signalling) and tumour-associated stress responses, including apoptosis regulation, response to stimuli and vascular development (Supplementary Fig. 14b). Cellular component and molecular function terms further highlighted extracellular matrix organisation, cell–cell communication and receptor binding (Supplementary Fig. 14a, Supplementary Fig. 15). Despite limited one-to-one correspondence with single-cell references, the *H* matrix recovers coherent and biologically meaningful axes of GBM variation, including lineage, hypoxia, inflammatory signalling and stromal remodelling.

## 3 Discussion

In this study, we introduce PISTACHIO, a proteomics-constrained framework for spatial transcriptomics deconvolution that integrates spatial protein information into a matrix factorisation model. By incorporating imaging mass cytometry (IMC)-derived spatial masks into a negative binomial NMF formulation, the method jointly estimates cell-type composition and transcriptional programs directly from spatial transcriptomics data without requiring matched scRNA-seq references.

Across synthetic and real datasets, our results show that incorporating proteomic spatial constraints substantially improves the recovery of spatial cell-type composition. In controlled simulations, PISTACHIO achieved near-perfect agreement with ground-truth and remained robust to substantial perturbations in the spatial prior, indicating that the model uses the mask as a guiding constraint rather than a rigid assignment. Compared with existing methods, PISTACHIO outperformed both reference-based approaches, such as Cell2location, and reference-free approaches, such as STdeconvolve, particularly in settings where single-cell references were mismatched or unavailable. These findings highlight the advantage of integrating orthogonal spatial information to reduce ambiguity in deconvolution.

Importantly, PISTACHIO also enables the recovery of cell-type-specific transcriptional programs through the inferred gene-loading matrix *H*. In prostate cancer, these programs showed clear correspondence with external single-cell references and aligned with known epithelial, stromal and immune compartments. In glioblastoma, where transcriptional states are highly plastic and cell identities form continua rather than discrete classes, direct one-to-one correspondence with single-cell references was weaker. However, functional enrichment analyses demonstrated that the inferred programs captured major axes of GBM biology, including hypoxia, inflammatory signalling, metabolic adaptation, mesenchymal transition and vascular remodelling. These results suggest that in highly heterogeneous tumours, functional coherence provides a more appropriate validation framework than strict matching to predefined cell-type labels.

A key feature of the proposed approach is the use of hard spatial constraints derived from proteomics. Unlike probabilistic priors used in existing models, the binary mask enforces spatial feasibility of cell-type presence while allowing abundance to be inferred from transcriptomic data. This design improves interpretability and prevents spurious assignment of cell-types to biologically implausible regions. At the same time, robustness analyses indicate that the model tolerates moderate inaccuracies in the mask, suggesting that it can accommodate realistic levels of noise arising from segmentation errors or imperfect co-registration between modalities.

The performance differences observed between prostate carcinoma and GBM further illustrate the impact of tissue architecture in deconvolution. In prostate tissue, which exhibits relatively well-defined glandular organisation, inferred cell-types and transcriptional programs align closely with both proteomic and transcriptomic references. In contrast, GBM represents a more challenging setting characterised by diffuse spatial organisation, overlapping transcriptional states and substantial tumour–microenvironment interactions. In this context, the ability of PISTACHIO to recover coherent functional programs despite incomplete single-cell correspondence highlights the value of integrating spatial priors and focusing on program-level interpretation.

Despite these strengths, several limitations should be considered. First, the spatial mask is treated as deterministic, whereas in practice IMC-derived annotations may contain uncertainty due to marker ambiguity, segmentation errors or registration mismatches. Extending the framework to incorporate probabilistic or learnable spatial constraints could improve flexibility in such settings. Second, evaluation of transcriptional programs in real datasets was limited by the lack of donor-matched scRNA-seq references. While functional enrichment provides indirect validation, future studies with matched multimodal datasets will enable more direct benchmarking of inferred gene-expression profiles. Finally, the current model assumes a fixed set of cell-types and does not explicitly model continuous cell-state transitions, which are prominent in tumours such as GBM.

Future work could extend PISTACHIO in several directions. Incorporating uncertainty-aware or hierarchical priors may allow the model to better capture continuous cell-state variation. Integration with additional modalities, such as spatial epigenomics or histology-derived features, could further refine spatial constraints and improve biological interpretability. More broadly, the framework provides a general strategy for embedding complementary spatial information into transcriptomic models, which may be applicable to a wide range of spatial multi-omics settings.

## 4 Methods

### 4.1 Synthetic data

We benchmarked PISTACHIO using synthetic spatial transcriptomics data generated with the Stereoscope framework (Andersson et al. 2020). Simulations were based on a harmonised glioblastoma multiforme (GBM) single-cell dataset (Ruiz-Moreno et al. 2022), selecting the donor with the largest number of cells (BT363). Spatial locations were constructed as mixtures of annotated cell-types, with proportions drawn from a Dirichlet distribution following the original Stereoscope implementation.

To better reflect sequencing-based spatial transcriptomics, we extended the Stereoscope simulation pipeline to track unique molecular identifiers (UMIs) at the single-cell level. For each simulated spot, selected cells were aggregated and expression counts were downscaled by a factor of 0.1 to Visium-like depth. This produced a spot-level count matrix *Y*, a depth matrix 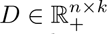 encoding ground-truth cell-type contributions, and a members matrix 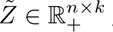 giving the number of cells of each type per spot. The binary mask used for constrained deconvolution was obtained as *Z* = I(*Z̃ >* 0). The final dataset comprised 2,450 spots, 9 cell-types, and 28,045 genes.

#### Cell-type abundance recovery

Model performance was assessed by comparing inferred cell-type abundances *W* to the ground-truth depth contributions *D*. We computed Pearson correlations between *W*_·*k*_ and *D*_·*c*_ across spots for all pairs of estimated components *k* and ground-truth cell-types *c*, excluding entries where both values were zero. Correlations were summarised in a cell-type-by-cell-type matrix, and agreement along the diagonal (matched labels) was aggregated by averaging.

#### Gene signature recovery

We assessed the recovery of gene expression signatures by comparing inferred profiles *H* to single-cell reference profiles computed as the mean scRNA-seq expression per cell type. To match preprocessing and model assumptions, scRNA-seq means were variance-scaled for Gaussian NMF models, whereas raw (unscaled) values were used for NB-NMF to preserve the negative binomial count model.

To focus signature comparisons on informative genes and reduce domination by ubiquitous housekeeping genes, we constructed marker-based gene sets from single-cell references. We obtained single-cell differential expression markers per cell-type using Scanpy’s `rank genes groups` (Wolf et al. 2018), selecting top markers ranked by adjusted *p*-values and forming a union across cell-types. As for *W* evaluation, we summarised diagonal (same-label) correlations by averaging.

#### Robustness to uncertainty in spatial priors

To evaluate robustness to uncertainty in spatial priors, we introduced structured perturbations to the binary cell-type presence mask derived by binarising the members matrix *Z̃*. Spatial coordinates were assigned on a synthetic two-dimensional grid to mimic Visium spot geometry and define local neighborhoods. We considered two biologically motivated noise models: (i) a local density model introducing density-dependent false positives and false negatives (e.g. reduced sensitivity in sparsely populated regions and increased ambiguity in densely packed areas), and (ii) a neighborhood switch model that partially replaces a spot’s mask with information from its local neighborhood, mimicking boundary effects and local mixing between adjacent populations (e.g. tumour/immune interfaces). We used *k* = 6 nearest neighbors and applied noise at levels *p* ∈ {0, 0.05, 0.10, 0.20}, with *p* = 0 corresponding to the noise-free setting. For each noise level and noise model, PISTACHIO was re-run using the perturbed spatial prior, and robustness was quantified by the averaged Pearson correlation between inferred and ground-truth cell-type contributions.

#### Unknown subtype recovery

To probe whether PISTACHIO can resolve hidden heterogeneity when prior information is coarsened, we designed an unknown subtype recovery experiment. We merged pairs of biologically related cell-types into a single “Unknown” category in the binary presence mask, while keeping spot-level expression counts unchanged. Two latent components were then assigned to the collapsed class, such that the prior no longer distinguished the original subtypes and the task reduced to blind source separation within this group. We evaluated three scenarios of increasing difficulty: (i) TAM-BDM versus TAM-MG (highly similar immune subtypes), (ii) Endothelial versus Mural cells (related but more transcriptionally distinct vascular types), and (iii) OPC-like versus NPC-like malignant programs (moderately similar tumour subpopulations). Performance was assessed using all-vs-all Pearson correlations between inferred components and ground-truth depth contributions.

### 4.2 Prostate cancer spatial multi-omics data

#### Human tissue samples

Human prostate samples were obtained from the Manchester Cancer Research Centre (MCRC) biobank under the biobank project 19 ROBR 01. This project was approved by the MCRC Biobank Research Tissue Bank Ethics (ref: 18/NW/0092). FFPE blocks were generated from the tumour regions of patients undergoing radical prostatectomy for localised prostate cancer. All samples were reviewed by a consultant genitourinary (GU) histopathologist after haematoxylin and eosin (H&E) staining. Samples with a minimum of 20% tumour content and of sufficient size for histopathological applications were selected for the study.

#### Spatial transcriptomics laboratory workflow

Tissue preparation and sectioning was performed in accordance with the Visium spatial gene expression for FFPE tissue preparation guide (10x Genomics, CG000408) and using a Visium spatial gene expression slide kit (10x Genomics, PN-1000188). Library construction and sequencing was performed according to the 10X Genomics user guide (CG000407, 10x Genomics), using the visium FFPE reagent kit (PN1000361) and human transcriptome probe kit (10x Genomics, PN-1000363). Firstly, probes were hybridised overnight followed probe ligation, probe release, probe extension and finally FFPE library construction. Sequencing was performed using a NovaSeq 6000 sequencer (Illumina) with paired-end 28+10+10+50 cycles and an SP flowcell with standard loading. Sequencing depth was calculated according to the sample coverage of the capture area such that a minimum of 25,000 paired-end reads were used per tissue covered capture area spot.

#### Imaging mass cytometry

Staining was performed according to Fluidigm instructions for FFPE tissues. Full details of antibody clones, metal labels and concentrations can be found in Supplementary Table 1. Antibodies not supplied by Fluidigm were conjugated using a Maxpar labelling kit (Fluidigm, USA), as per manufacturer’s instructions. Final concentrations of conjugated antibodies were determined by Nanodrop (Thermofisher), before being reconstituted in Antibody Stabilisation Solution (Candor Bioscience, Germany), and stored at 4°C. IMC images of tissue sections bound with metal-conjugated antibodies were acquired using a Hyperion imaging mass cytometer (Fluidigm). In brief, the tissue was laser-ablated in a rastered pattern in a series of 1 µm2 pixels. The resulting plume of ablated tissue was then passed through a plasma source, ionising it completely into its constituent atoms. Time-of-flight mass spectrometry then discriminated the signal for each of the metal-conjugated antibodies, and images for each antibody were reconstructed based off the metal abundancy at each pixel. Multi-channel images for each were exported from the raw data as OME-TIFFs using MCD Viewer (Fluidigm).

#### Spatial transcriptomics data pre-processing, quality check and filtering

Pre-processing of Visium data was performed by submitting fastq files along with full-resolution TIFF images to the Space Ranger count pipeline (version 1.3.0) using the ‘–reorient-images’ flag and GRCh38 reference; QC metrics for spots were assessed using Space Ranger report outputs. Data filtration for lowly expressing genes and spots was performed by removing spots with fewer than 100 unique genes expressed, spots with less than 500 unique molecular identifier (UMI) count, and genes expressed in less than 5 spots. Raw count normalisation was performed with SCTransform33 (SCT) using the top 3000 variable features.

Further analyses were performed for two samples (ST2 and ST1). Downstream Visium handling used scanpy/anndata objects containing the raw spot-by-gene count matrix *Y* and spot coordinates. For sample ST2, a subset of spatial spots was manually excluded due to a visually identified misread tissue region. Briefly, the high-resolution histology image (hires) and spot coordinates were extracted from the Visium AnnData object and mapped to image pixel space using the Space Ranger high-resolution scale factor. Spots were overlaid on the histology image and a lasso tool (matplotlib.widgets.LassoSelector) was used to interactively select the misread region. Selected spot barcodes were stored in adata.obs["misread region"] and exported to misread spots barcodes.csv. A filtered Visium object (adata clean) was created by removing these spots prior to multimodal alignment and deconvolution.

#### IMC preprocessing, clustering, and rule-based cell-type annotation

IMC single-cell measurements were exported from HALO as per-cell feature tables and loaded into pandas (Indica Labs). Duplicate objects (based on identical bounding-box coordinates: XMin, XMax, YMin, YMax) were removed. We retained protein channels corresponding to marker intensities (columns containing "Cell Intensity") and excluded DNA channels (Ir191 and Ir193). Marker intensities were transformed using an arcsinh transformation with cofactor 5, scaled, reduced by principal component analysis (PCA), and a *k*-nearest-neighbour graph was constructed in the PCA space using scanpy. Cells were clustered with the Leiden algorithm, using resolution 0.4 for ST2 and 0.5 for ST1.

To assign biological cell-type labels, we computed cluster-level mean expression for a curated prostate IMC marker panel covering epithelial/tumour, immune, and stromal populations. For each Leiden cluster, a score was computed for each candidate label as the mean intensity across its associated marker channels, and the highest-scoring label was assigned. To improve separation of T cell subsets, we applied a rule-based refinement using CD3, CD4, and CD8 marker expression: clusters with high CD3 and high CD8 were labelled as CD8 T; clusters with high CD3 and high CD4 were labelled as CD4 T; otherwise the highest-scoring label was used. UMAP visualisation was used for qualitative inspection of clustering and annotations.

#### Co-registration of Visium Spot Loci

IMC and H&E TIFF images from serial sections were co-registered in HALO using b-spline transformation and manual landmarking. Visium spot loci, relative to full-res H&E, were extracted from space ranger outputs and converted to HALO-compatible XML annotation files. Annotations were imported into HALO and spot loci transferred to IMC-relative loci via the co-registration transform.

#### Spot-level IMC ground-truth and spatial priors

IMC cell-to-spot membership was derived from the HALO `"Analysis Region"` field, which encodes Visium spot barcodes. For sample ST2, IMC rows associated with the manually excluded Visium barcodes (from `misread spots barcodes.csv`) were removed prior to constructing spot summaries. Spot-level IMC-derived cell counts were computed as a contingency table *Z̃* ∈ ℝ*^n^*^×*k*^ using `pandas.crosstab`, where *n* is the number of spots and *k* is the number of IMC-defined cell-types. A corresponding binary presence mask *Z* = I(*Z̃ >* 0) was constructed and used as a spatial prior for constrained deconvolution. In addition to cell counts, we also computed an area-weighted ground-truth by summing per-cell areas within each spot and cell-type (HALO column `Cell Area` (*µ*m^2^)), producing a spot-by-type matrix of total occupied area. For comparisons of compositions, both the estimated abundances and the area-based ground-truth were optionally row-normalised to sum to one per spot.

#### Biologically motivated coarse cell classes

Fine-grained IMC annotations were additionally grouped into biologically interpretable coarse classes for deconvolution and evaluation using a predefined mapping. Specifically, epithelial-related subtypes (Epithelial, Proliferative Tumor, Hypoxic Tumor, Stem-like/Basal) were grouped into an epithelial super-class; T cell and cytotoxic subtypes (CD4 T, CD8 T, T näıve/memory, NK-like cytotoxic) were grouped into a T cell super-class; myeloid-related subtypes (Myeloid/Macrophage, M2-like TAM, Myeloid APC) into a myeloid super-class; CAF/Myofibroblast into a stromal myofibroblast class; and B cells were retained as a separate class. This hierarchy was applied both to IMC ground-truth (counts and area) and, where needed, to deconvolution outputs by summing estimated abundances across fine types that map to the same coarse class.

#### Alignment of IMC and Visium

To align modalities, Visium spots were restricted to those with IMC correspondence based on shared barcodes between the filtered Visium object and the IMC-derived spot tables. Gene names in the Visium object were made unique before subsetting. After alignment, the Visium count matrix *Y* ∈ ℝ*^n^*^×*g*^ was extracted (dense if required) and used as input for NMF-based deconvolution, with cell-type abundances represented by *W* ∈ ℝ*^n^*^×*k*^ and gene-expression signatures by *H* ∈ ℝ*^k^*^×*g*^, where *k* corresponds to the set of deconvolved cell-types/classes and *g* is the number of genes.

#### Constrained negative binomial NMF deconvolution

Deconvolution was performed using a negative binomial NMF objective (implemented in our code as `nmf_main`), using the binary presence mask *Z* as a constraint on spot–cell-type support. The method returns factor matrices *W* and *H*, the optimisation trace (training loss across iterations), and reconstructed counts *Ŷ* = *WH*. Model fit was assessed by comparing *Y* and *Ŷ* using global metrics (e.g. *R*^2^, MAE, Pearson correlation on vectorised entries) and by visualising the training curve and *Y* versus *Ŷ* scatter plots.

#### Evaluation against IMC-derived ground-truth

We evaluated inferred abundances *W* against IMC-derived spot-level ground-truth using Pearson correlation. Evaluations were performed both at the fine label level (spot-by-fine-type) and after collapsing to the biological coarse classes described above. In addition to count-based ground-truth, we evaluated against the area-based ground-truth derived from summed cell areas per spot and type. Correlation matrices (estimated types versus ground-truth types) were computed after aligning spots, optionally dropping spot entries where both estimated and ground-truth values were near zero. Summary metrics included row-wise best-match correlations and same-name (diagonal) correlations where label sets matched.

#### Validation of inferred gene-expression signatures using scRNA-seq

Inferred signatures *H* were validated against an external single-cell prostate cancer atlas. Gene symbols were harmonised by uppercasing and intersecting the atlas gene symbols with the Visium/NMF signature gene set. The scRNA-seq data were total-count normalised (target sum 10^4^) and log-transformed. Marker genes per scRNA-seq cell-type were identified using a Wilcoxon rank-sum test (`scanpy.tl.rank_genes_groups`). To evaluate agreement between reference and inferred signatures, we computed cell-type mean expression profiles from scRNA-seq and compared them to *H* over shared genes using three similarity measures: Pearson correlation, Spearman correlation, and cosine similarity (after per-gene standardisation). For each similarity matrix, we summarised performance using row-wise best matches and additionally computed a one-to-one assignment between scRNA-seq types and inferred types using the Hungarian algorithm, enabling matched scatter plots of scRNA-seq mean expression versus inferred signature values for the top matched pairs.

#### Pathology-aware gene panel decomposition

To interpret deconvolution results in the context of tissue pathology, we generated per-gene spatial panels that juxtapose observed Visium expression with model reconstructions and cell-type contributions. For each sample, we loaded the spot-level cell-type abundance matrix *W* and the corresponding gene-loading matrix *H*, aligned them to shared barcodes, and restricted *H* to genes present in the Visium feature space. For each selected pathology-associated gene *j*, we extracted the observed spot-level expression vector *y*_·*j*_ from the Visium count matrix (optionally from a specified layer). We computed the reconstructed expression as *ŷ*_·*j*_ = *Wh_j_*, where *h_j_* denotes the column of *H* for gene *j*. Spot-wise cell-type contributions were calculated as *c*_·*jk*_ = *W*_·*k*_*H_kj_*, which sum to *ŷ*_·*j*_ across cell-types. When *W* represented row-normalised abundances, we rescaled *ŷ* and *c* to the observed sequencing depth by matching the predicted total expression per spot to the observed library size (row sum of the Visium expression matrix), ensuring comparability of reconstructed and observed counts. As an alternative view, we optionally reported all quantities as per-spot proportions by dividing *y*, *ŷ*, and *c* by the library size to mitigate depth effects.

For visualisation, we selected the top *K* contributing cell-types for each gene based on their global contribution share *_i_ c_ijk_/ _i_ ŷ_ij_* and displayed a single-row multi-panel figure comprising (i) the categorical histology annotation, (ii) observed expression *y*_·*j*_, (iii) reconstructed expression *ŷ*_·*j*_, and (iv) the top-*K* contribution maps *c*_·*jk*_ (optionally including a residual map *y*_·*j*_ − *ŷ*_·*j*_). Continuous panels were optionally log-transformed using log(1 + *x*) to stabilise dynamic range and were clipped using shared percentile bounds (default 1st–99th) to reduce the influence of extreme hotspots. All continuous panels used a shared colour scale and a single colourbar for direct visual comparison, with optional cropping to a user-defined spatial region of interest (ROI) in Visium coordinates.

#### Construction of prostate gene programs

To interpret spatial transcriptomic patterns in the prostate tumour microenvironment, we constructed a curated set of prostate-specific gene programs capturing epithelial identity, tumour biology, and stromal and immune signalling (reviewed in (Ali et al. 2026)). Candidate genes were compiled through systematic review of prostate spatial transcriptomics, single-cell atlases, and molecular characterization studies of prostate cancer (Mikutenaite et al. 2025; Suvac et al. 2025; Chen et al. 2021; Dutta et al. 2025; Krossa et al. 2025; Liu 2024; Attard et al. 2016; Bernardini et al. 2026; Miao et al. 2021; Gheybi et al. 2025; Ospina et al. 2024; Desai et al. 2024; Wang et al. 2025; Shayit et al. 2025; Li et al. 2025; Zhong et al. 2025; Fonteyne et al. 2026).

#### Program scoring and comparative visualization

To compare transcriptional programs between samples, we computed program scores using the gene loadings matrix *H* obtained from the constrained matrix factorization model. For each sample separately, gene weights in *H* were standardised across cell-types using gene-wise z-score normalisation to account for differences in scale across genes. Specifically, for each gene *j*, the standardised value was computed as

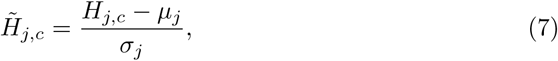

where *H_j,c_*denotes the gene loading for gene *j* in cell-type *c*, and *µ_j_* and *σ_j_* represent the mean and standard deviation of that gene across cell-types within the sample.

Gene programs were defined using curated gene sets (see `Construction of prostate gene programs`). For each cell-type and program, a program score was calculated as the mean z-score of genes belonging to that program, considering only genes present in the *H* matrix and requiring a minimum of two genes per program. This resulted in a program-by-cell-type score matrix for each sample.

To compare programs between samples (sample ST1 vs. sample ST2), we computed the difference in mean program scores for each cell type:

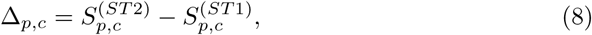

where *S_p,c_*represents the mean z-score for program *p* in cell-type *c*.

Statistical significance of program differences was assessed using a gene-level bootstrap sign-flipping test. For each program, the difference in gene-level z-scores between samples was calculated for all genes belonging to that program. Under the null hypothesis of no systematic difference, the sign of each gene-level difference was randomly flipped across bootstrap iterations (5,000 iterations). The bootstrap distribution of mean differences was used to compute two-sided *p*-values. *P* -values were subsequently adjusted for multiple testing within each cell-type using the Benjamini–Hochberg false discovery rate (FDR) procedure.

Results were visualized using point plots, where each point represents the mean program score for a given cell-type and sample. Programs were displayed along the x-axis and mean gene z-scores along the y-axis, with samples distinguished by colour. Statistical significance of the difference between samples was indicated using standard significance annotations based on FDR-adjusted *p*-values.

### 4.3 Glioblastoma spatial multi-omics data

#### Human tissue samples

Formalin-fixed paraffin-embedded (FFPE) tissue sections from four primary IDH wild-type glioblastoma (GBM) cases were obtained from the Department of Cellular Pathology at Salford Royal Hospital under University of Manchester ethics (Integrated Research Application System (IRAS) ID 244538, reference: 19/NE/0186). Informed consent was obtained from patients for use of their tissue for research. Tissue blocks were sectioned at 5 µm thickness and mounted onto Visium Spatial Gene Expression slides. Sections were stored under appropriate conditions prior to processing, following manufacturer protocols.

#### Spatial transcriptomics workflow

Tissue sections were deparaffinised in xylene, and rehydrated through a graded ethanols according to the 10x Genomics protocol. A section of each case was stained with haematoxylin-eosin to be used as a reference to interpret the transcriptomics and proteomics data. The slides were acquired at high resolution with a Pannoramic brightfield scanner (3DHISTECH).

Spatial gene expression profiling was performed using the Visium Spatial Gene Expression for FFPE workflow (10x Genomics). Briefly, gene-specific probe pairs were hybridised to RNA targets in situ, followed by probe ligation and release. Captured probe products were amplified to generate spatially barcoded sequencing libraries according to the manufacturer’s instructions. Libraries were quality controlled and sequenced on an Illumina NextSeq 500 system at the University of Manchester Genomics Facility. Sequencing reads were processed and mapped to spatial coordinates using Space Ranger, and downstream visualisation was performed using Loupe Browser.

Due to suboptimal H&E staining on the Visium tissue section, the images of the serial H&E-stained sections were co-registered to the Visium capture area to enable spatial alignment of gene expression data with tissue morphology.

#### Imaging mass cytometry acquisition and preprocessing

Adjacent FFPE sections were profiled using Hyperion imaging mass cytometry (Standard BioTools). Tissue sections (5 µm thickness) were deparaffinised and subjected to antigen retrieval at 96^◦^C for 30 minutes in Tris-EDTA (pH 8.5). Non-specific binding was blocked using 3% bovine serum albumin for 45 minutes, followed by overnight incubation at 4^◦^C with lanthanide-conjugated antibodies diluted in PBS with 0.5% BSA. Antibodies were conjugated using Maxpar Antibody Labeling Kits (Standard BioTools).

Slides were washed with PBS and 0.1% Triton-X100, stained with iridium (1:400, Intercalator-Ir, Standard BioTools) for nuclear labelling, rinsed with ultrapure water, and air-dried. Images were acquired using a Hyperion imaging mass cytometer, where tissue was laser-ablated in a rastered pattern at 1 µm^2^ resolution. The resulting plume was ionised in a plasma source, and time-of-flight mass spectrometry was used to quantify metal-tagged antibodies, reconstructing multiplexed protein images. Staining quality was reviewed using MCD Viewer by a pathologist.

IMC images were denoised using the IMC-Denoise algorithm (Lu et al. 2023). Cell segmentation was performed using CellPose 2.0 (Pachitariu and Stringer 2022), generating cell masks that were applied to all antibody channels to obtain single-cell protein expression profiles. Marker intensities were normalised to their 99th percentile values.

Single-cell IMC data were analysed in Python using scanpy (Wolf et al. 2018). Dimensionality reduction and clustering were performed using the Leiden algorithm (Traag et al. 2019), and clusters were manually annotated into tumour, immune, and stromal populations based on established marker expression profiles.

#### Spatial alignment and construction of multimodal matrices

IMC cell centroids were co-registered to Visium spatial barcodes, restricting analysis to matched spatial locations. This produced a spot-level IMC-derived cell count matrix *Z̃* ∈ ℝ*^n^*^×*k*^, where *n* denotes the number of spatial locations and *k* the number of IMC-defined cell-types, alongside a corresponding binary presence mask *Z* = I(*Z̃ >* 0). The matched Visium gene-expression matrix is denoted as *Y* ∈ ℝ*^n^*^×*g*^.

The GBM cohort comprised four samples (A–D). Sample A (N540) contained 2,849 spatial locations and 17,943 genes with 26 IMC-defined cell-types. Sample B (N935) contained 1,656 spatial locations and 17,943 genes with 24 cell-types. Sample C (N403) contained 4,115 spatial locations and 17,943 genes with 24 cell-types. Sample D contained 1,656 spatial locations and 17,943 genes with 26 cell-types. After filtering to genes retained for deconvolution, the corresponding gene-loading matrices *H* had dimensions 26 × 11,842 (A), 24 × 11,875 (B), 24 × 11,784 (C), and 26 × 11,308 (D).

#### Cell-type aggregation and biological interpretation

IMC annotations comprised a heterogeneous set of tumour, immune, and stromal cell states, including rare and transitional populations. To improve robustness and interpretability, fine-grained cell-type labels were grouped into broader, biologically meaningful categories reflecting established GBM transcriptional programs (Neftel et al. 2019). Tumour cells were aggregated into astrocyte-like, OPC-like, NPC-like, and mesenchymal-like lineages, while immune populations were grouped into a myeloid compartment and non-malignant cells into a vascular/stromal compartment. Spot-level abundances for these classes were obtained by summing the corresponding entries in *Z̃*.

#### Deconvolution and evaluation

The matrices *Y*, *Z̃*, and *Z* were used as inputs and spatial priors for the PISTACHIO framework, as described for the prostate analysis. Because matched scRNA-seq ground-truth was not available for these FFPE GBM samples, evaluation focused on comparing inferred cell-type abundances *W* with IMC-derived spatial compositions.

Performance was assessed both quantitatively, via agreement with IMC-derived spot-level compositions, and qualitatively, through consistency with known GBM tumour microenvironment organisation. The pronounced inter-patient transcriptional heterogeneity characteristic of GBM (Ruiz-Moreno et al. 2022) limits direct evaluation of inferred gene-expression signatures *H*, and evaluation therefore emphasised spatial localisation patterns and robustness of inferred cell-type distributions.

### 4.4 Comparison with existing tools

To benchmark our NMF-based framework against existing deconvolution approaches, we applied STdeconvolve, an unsupervised probabilistic method based on Latent Dirichlet Allocation (LDA) that infers latent topics corresponding to putative cell-types (Miller et al. 2022). Following the recommended workflow, we filtered low-quality genes and low-coverage spots and selected highly overdispersed genes prior to model fitting to reduce noise and improve topic identifiability. LDA models were trained across a range of topic numbers *K*, and the optimal model was selected in a data-driven manner using STdeconvolve’s model selection criteria rather than being fixed *a priori*. This resulted in dataset-specific optimal topic numbers (*K* = 9 for the synthetic dataset, *K* = 6 for glioblastoma, and *K* = 5 for prostate). The selected model yields a topic–spot distribution (*θ*) and a topic–gene matrix (*β*), interpreted as inferred cell-type proportions and expression signatures, respectively. We evaluated *θ* by computing pairwise Pearson correlations with the ground-truth cell-type composition matrix *Z̃* (for the synthetic data) and assessed *β* by correlating topic-specific gene programs with reference mean gene expression profiles from single-cell data.

We next benchmarked our approach against Cell2location, a supervised Bayesian deconvolution method that leverages scRNA-seq reference profiles. For synthetic data, Cell2location was run using the matched single-cell reference (BT363). To assess robustness to inter-donor variation, we additionally used references from two other GBM donors (G967 and BT397) as well as a pan-donor reference. Donor BT397 shared the same tumour subtype as BT363, whereas donor G967 displayed broader heterogeneity (4/12 subtypes differed), enabling evaluation under biologically realistic reference variability. Cell2location was run with recommended settings unless otherwise specified, and with modified hyperparameters incorporating the number of cells per spot via the `N_cells_per_location` prior to test the impact of proteomics-derived constraints. Cell-type abundance estimates were compared against IMC-derived ground-truth for synthetic and real cancer data (prostate and GBM datasets), where IMC cell-type counts and matched scRNA-seq profiles served as informative priors.

## Supplementary information

Supplementary information is provided as a separate PDF.

## Acknowledgements

This work was supported by the European Laboratory for Learning and Intelligent Systems (ELLIS). E.B.I. acknowledges the ELLIS PhD Programme for supporting her doctoral research. We thank the Bioinformatics and Systems Biology group at the University of Manchester for helpful discussions and for providing access to the glioblastoma and prostate cancer datasets used in this study. We thank the Flow Cytometry and the Bioimaging Core Facilities at the University of Manchester for helping with the imaging mass cytometry work performed in this study. We thank the Genomics Technologies Core Facility at the University of Manchester for helping with the Visium ST work performed in this study.

## Declarations

### Ethics approval and consent to participate

Human prostate tissue samples were obtained from the Manchester Cancer Research Centre (MCRC) Biobank under project 19 ROBR 01, approved by the MCRC Biobank Research Tissue Bank Ethics Committee (reference: 18/NW/0092). Glioblastoma tissue samples were obtained from the Salford Royal Hospital under University of Manchester sponsored ethics (IRAS ID: 244538; reference 19/NE/0186). All samples were collected with patients’ informed consent and handled in accordance with the University of Manchester research governance framework.

### Consent for publication

Not applicable.

### Data availability

The spatial transcriptomics and imaging mass cytometry datasets analyzed herein were provided by the University of Manchester. The datasets generated during this study will be made publicly available upon publication.

### Materials availability

Not applicable.

### Code availability

The PISTACHIO software implementation and associated documentation are available as open-source software at https://github.com/ManchesterBioinference/PISTACHIO.

### Author contributions

E.B.I. conceived and implemented the PISTACHIO framework, performed all analyses, and drafted the manuscript. M.A.A., S.G. and M.R. conceived and supervised the study. M.J.H., L.B., A.H.A., F.R., K.P.H., K.C., D.W., R.S., P.O., A.B, J.A., and R.B. contributed to data generation, curation, and interpretation. All authors contributed to discussion of results and manuscript revision and approved the final version.

