## Supplementary Material for "Proteomics-constrained deconvolution reveals spatial cell-type programs in tumours"

Isik et al.

June 22, 2026

### S1 Supplementary Table

Table 1: Antibodies used for Hyperion imaging mass cytometry

| Antibody | Supplier | Product code | Dilution | Channel (mass tag) |
|---|---|---|---|---|
| SMA | Bio-Rad | MCA5781GA | 1:50 | 89 |
| PanCK | Biolegend | 914201 | 1:50 | 139 |
| CD11c | Abcam | ab52632 | 1:100 | 154 |
| HIF1$\alpha$ | Abcam | ab51608 | 1:100 | 155 |
| CD4 | Fluidigm | 91H010 | 1:50 | 156 |
| CD68 | Fluidigm | 91H012144 | 1:100 | 159 |
| CD8 | Fluidigm | 91H015 | 1:50 | 162 |
| GLUT1 | Abcam | ab115730 | 1:100 | 163 |
| $\beta$-Catenin | Fluidigm | 3165027A | 1:100 | 165 |
| CD74 | Fluidigm | 3166025D | 1:100 | 166 |
| CD45RO | Biolegend | 304202 | 1:100 | 173 |
| HLA-DR | Abcam | ab20181 | 1:100 | 174 |
| CD206 | CST | 91992 | 1:100 | 175 |
| MCT4 | Proteintech | 22787-1-AP | 1:100 | 176 |

### S2 Supplementary Figures

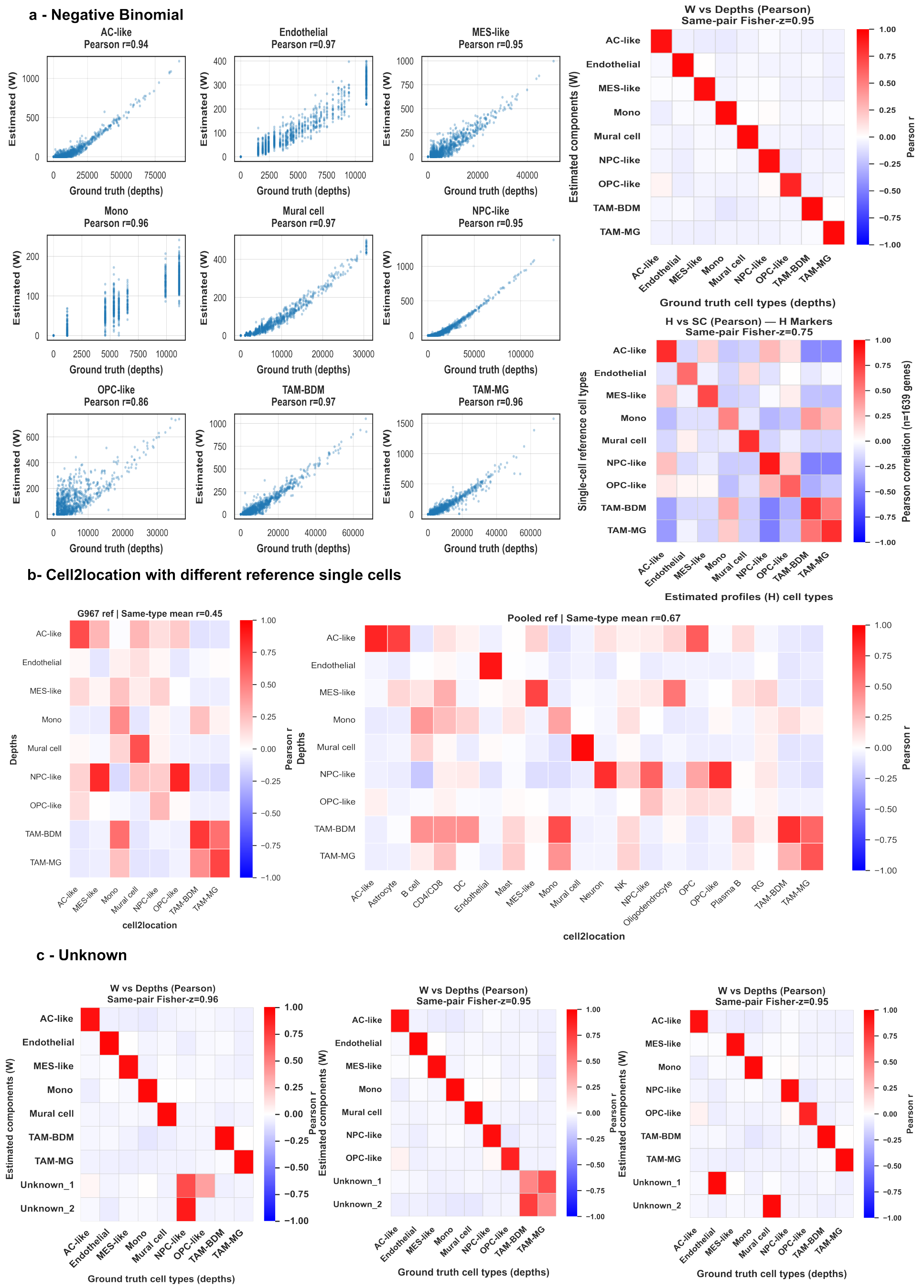


**Supplementary Figure S1: Additional synthetic benchmarking analyses.** **(a)** Detailed results for NB-NMF (PISTACHIO). (i) Per-cell-type scatter plots comparing inferred abundances $W$ with ground-truth depths across spatial spots. (ii) Pearson correlation heatmap between inferred abundances and ground-truth depths excluding shared zero entries. (iii) Pearson correlation between inferred gene-expression programs $H$ and single-cell reference signatures using marker genes derived from the estimated $H$ profiles. **(b)** Sensitivity of Cell2location to reference choice. Pearson correlation heatmaps between inferred abundances and ground truth when using a different-subtype reference and a pan-donor reference. **(c)** Unknown subtype recovery under coarsened priors. Pearson correlation between inferred components and ground-truth depths when two related cell types are merged into a single “Unknown” category in the spatial prior and two components are assigned to the collapsed group.

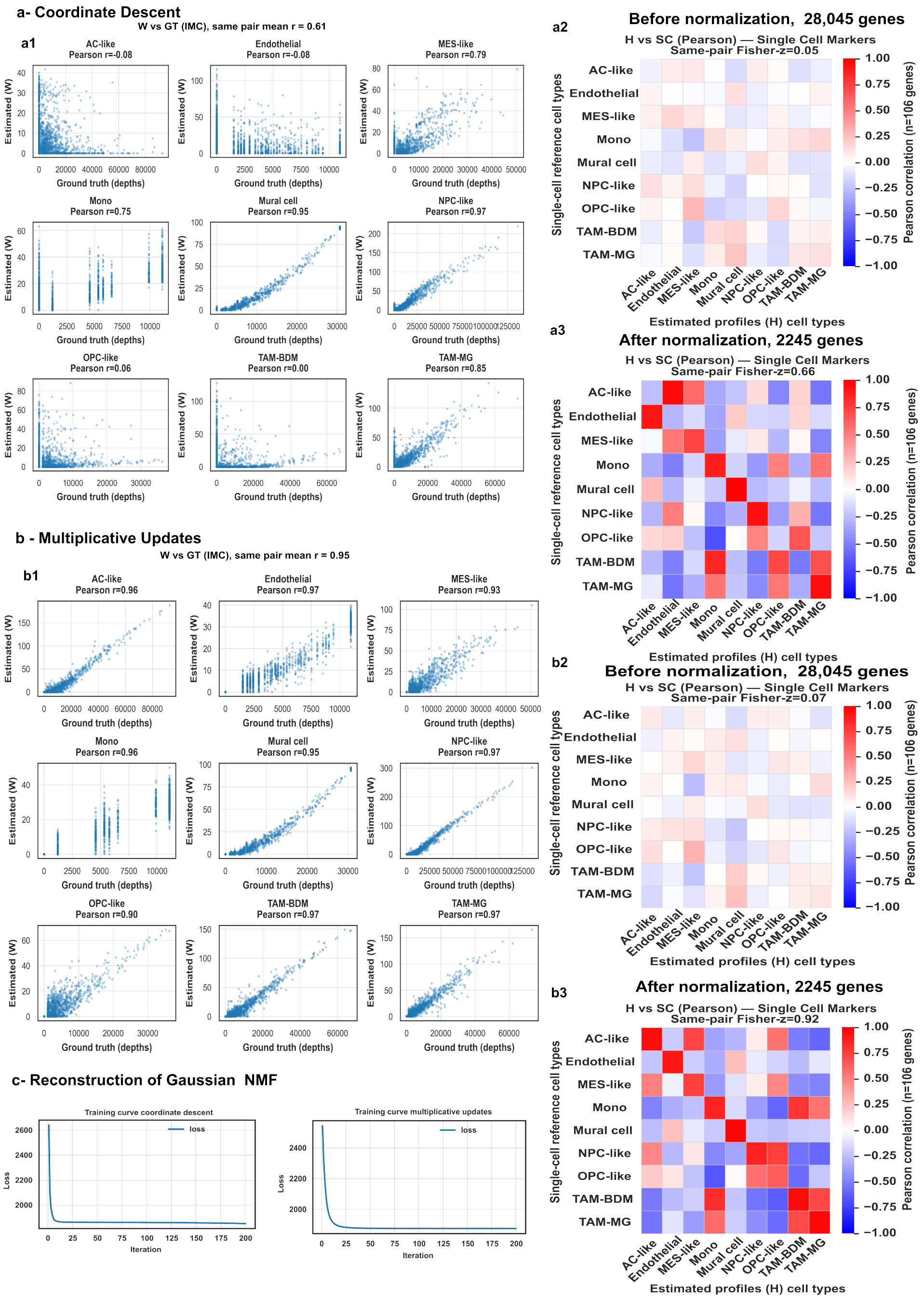


**Supplementary Figure S2: Comparison of Gaussian and negative binomial NMF formulations on synthetic data.** **(a)** Gaussian NMF optimised using coordinate descent. Left, per-cell-type scatter plots comparing inferred abundances $W$ with ground-truth depths. Right, Pearson correlation between inferred gene-expression programs $H$ and single-cell reference signatures before and after normalisation. **(b)** Gaussian NMF optimised using multiplicative updates. Left, per-cell-type scatter plots comparing inferred abundances with ground truth. Right, correlation between inferred gene programs and single-cell reference signatures before and after normalisation. **(c)** Reconstruction loss during optimisation for Gaussian NMF variants.

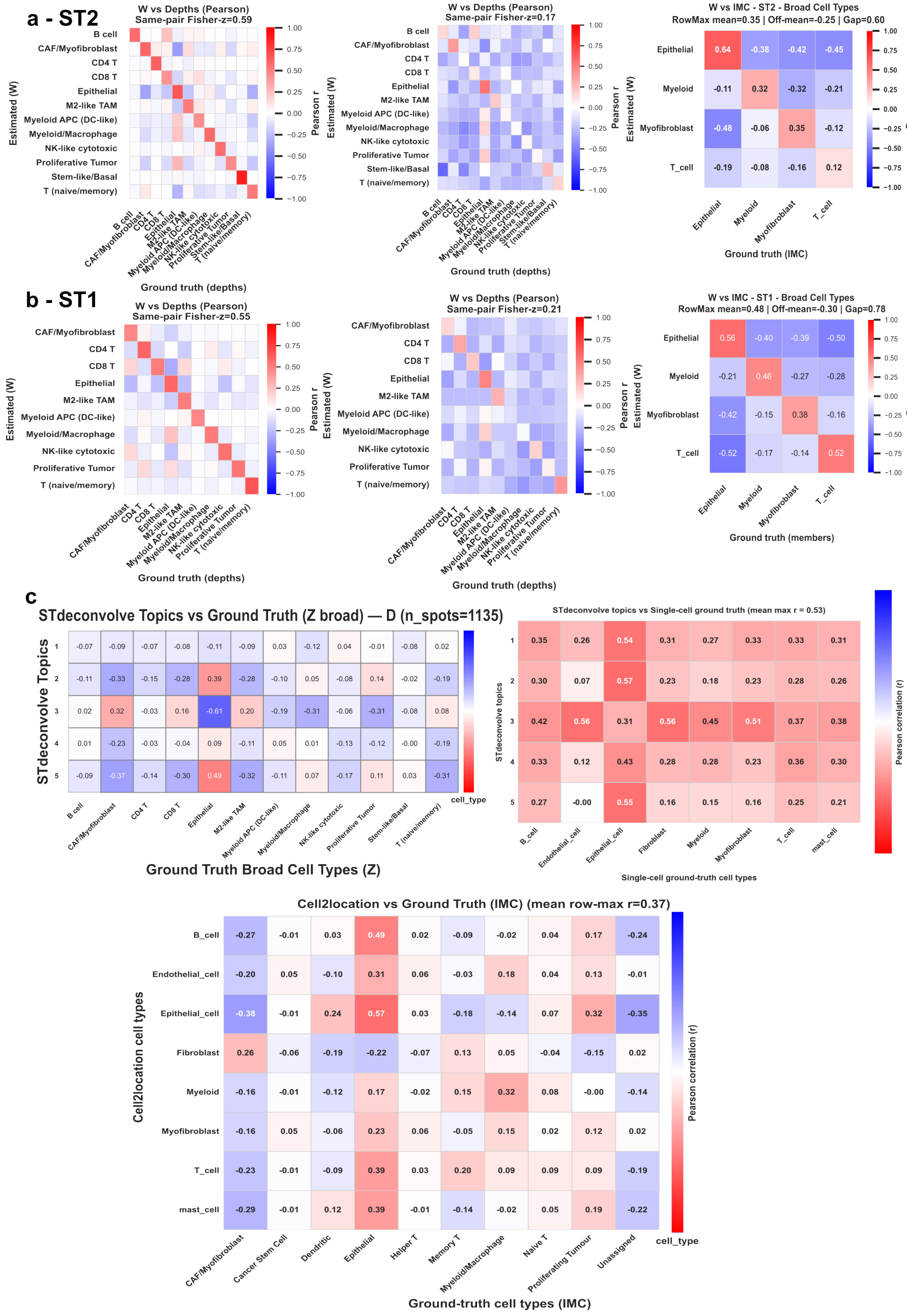


**Supplementary Figure S3: Additional benchmarking against IMC-derived ground truth for prostate samples.** **(a)** Pearson correlation between PISTACHIO-inferred cell-type proportions ($W$) and IMC-derived ground truth for prostate section ST2. Heatmaps show correlations computed using all spatial spots including shared zero entries (left), excluding shared zero entries (middle), and correlations aggregated across broad cell-type categories (right). **(b)** Corresponding correlation analyses for prostate section ST1, shown as in (a). **(c)** Comparison with alternative deconvolution methods. Top-left, correlation between STdeconvolve topics and IMC-defined cell types across spatial spots. Top-right, correlation between STdeconvolve topics and reference single-cell RNA-seq cell types using marker genes. Bottom, correlation between Cell2location-inferred cell types and IMC-derived ground truth. Colour scales indicate Pearson correlation coefficients.

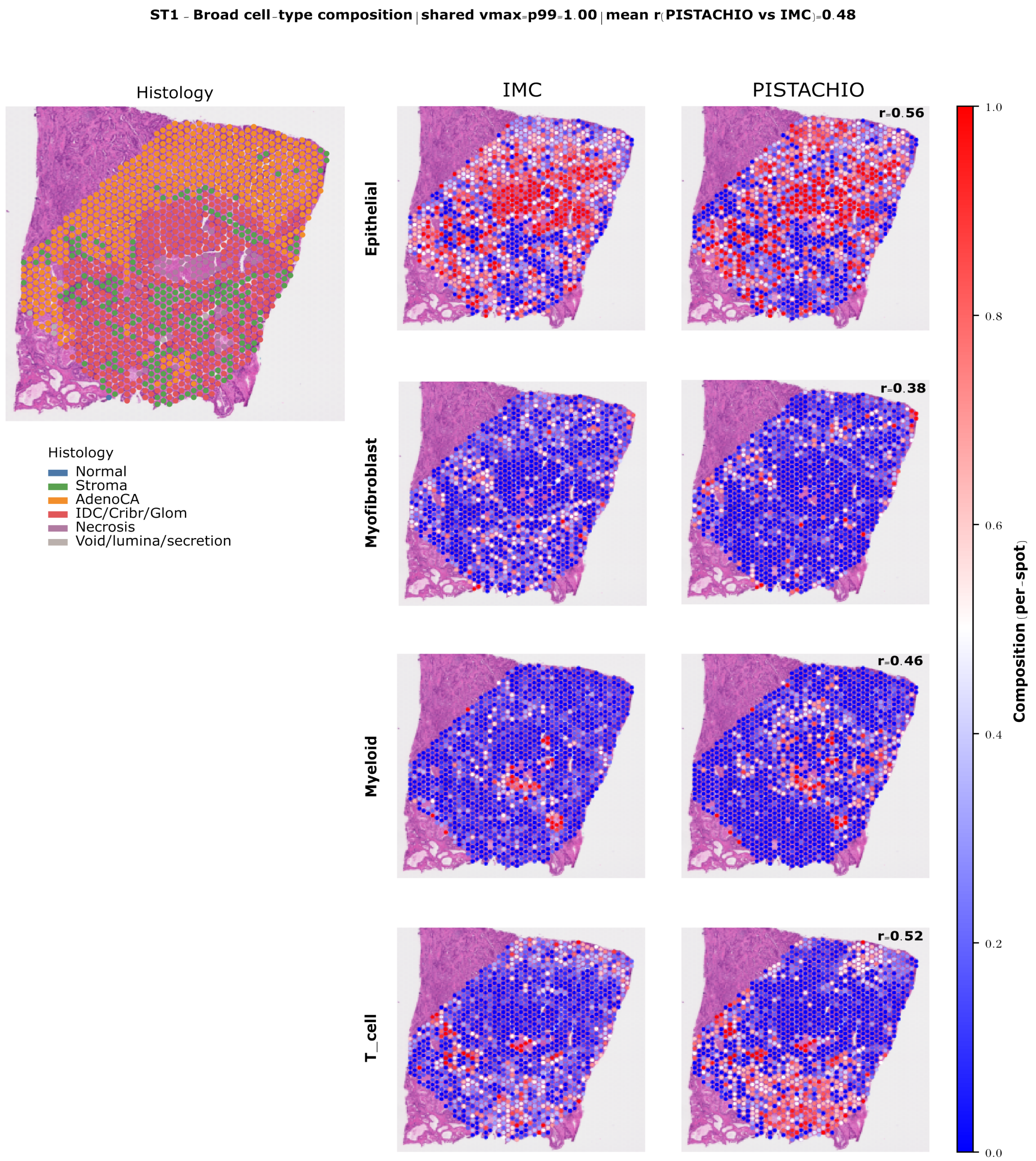


**Supplementary Figure S4: Spatial cell-type composition in prostate section ST1.** Histological annotation, IMC-derived ground truth, and PISTACHIO-inferred spatial composition are shown for four major cell-type compartments. Left, histology-based tissue annotation highlighting normal tissue, stromal regions, adenocarcinoma (AdenoCA), IDC/cribriform/glomeruloid patterns, necrosis, and luminal/secretory spaces. Middle, IMC-derived spatial proportions for epithelial, myofibroblast, myeloid, and T cell populations across Visium spots. Right, corresponding cell-type compositions inferred by PISTACHIO from spatial transcriptomics data. Pearson correlation coefficients between inferred and IMC-derived compositions are indicated for each cell type. Colours represent per-spot cell-type proportions using a shared colour scale across panels.

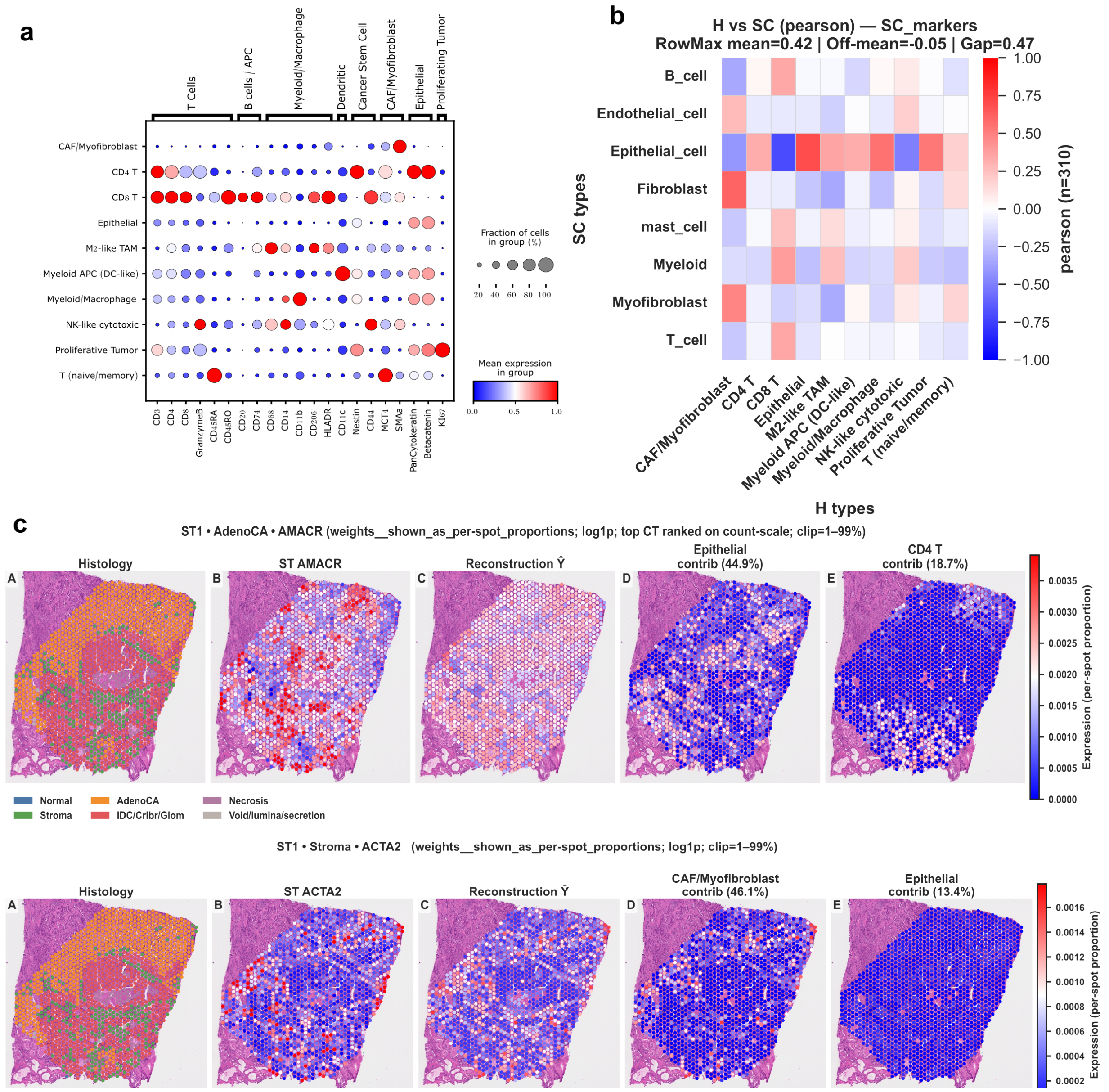


**Supplementary Figure S5: Validation of inferred transcriptional programs and gene-level spatial reconstruction in prostate section ST1.** **(a)** IMC protein marker profiles used to define fine-grained cell types for spatial constraints. Dot size indicates the fraction of cells expressing each marker, and colour indicates mean protein expression. **(b)** Correspondence between inferred transcriptional programs ($H$) and external single-cell RNA-seq reference cell types based on curated marker genes. The heatmap shows Pearson correlation between inferred programs and reference cell-type profiles. **(c, top)** Spatial reconstruction of the tumour-associated gene *AMACR*. Columns show histological annotation, observed spatial transcriptomics signal, reconstructed expression $\hat{Y}$, and dominant contributing cell types. **(c, bottom)** Spatial reconstruction of the stromal marker *ACTA2* in the same tissue section. Expression values are shown as per-spot expression levels after log transformation and clipping (1–99%).

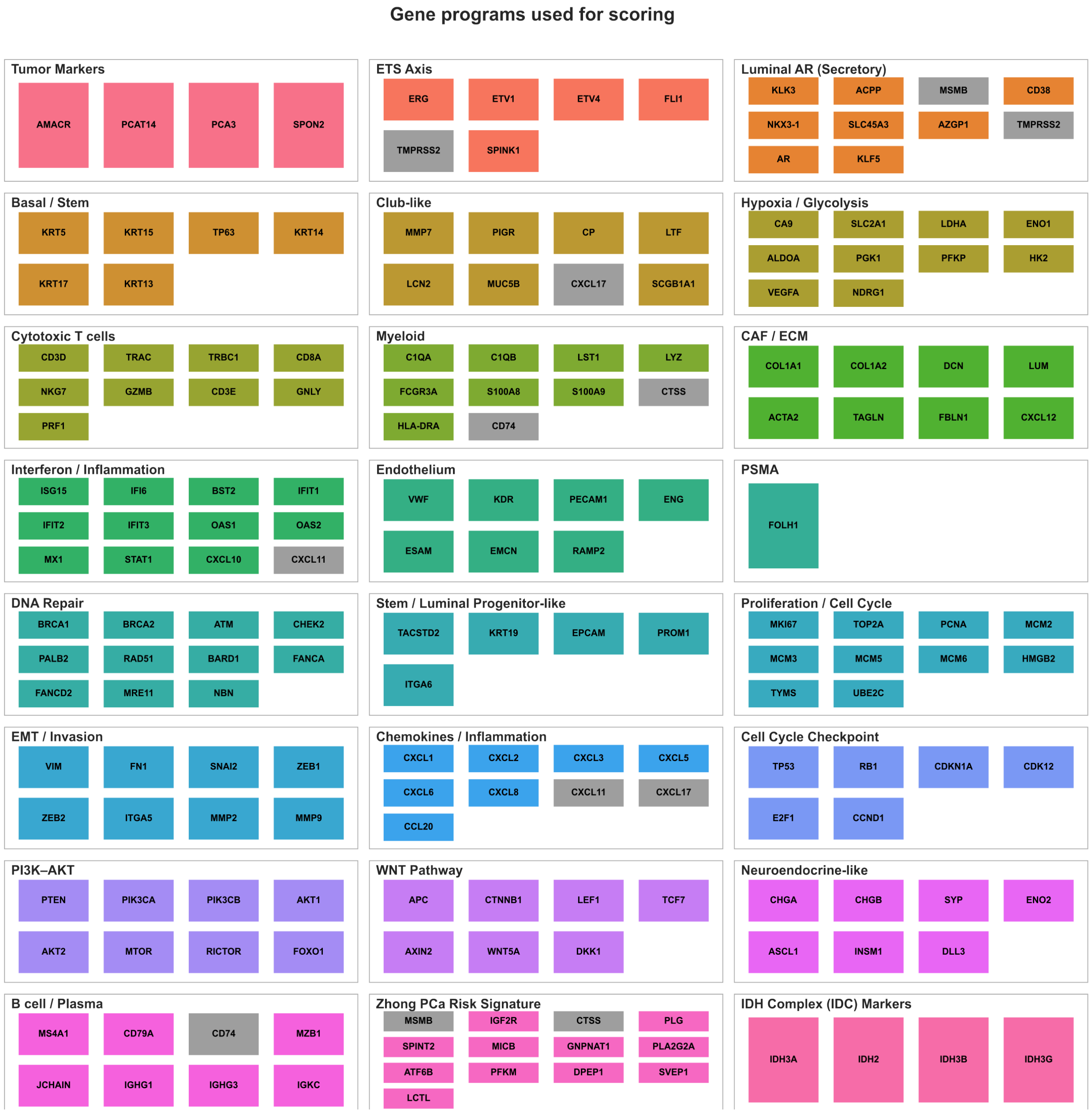


**Supplementary Figure S6: Curated gene programs used for transcriptional scoring.** Curated gene sets used to define transcriptional programs for program-level analyses across prostate tumour samples. Grey boxes indicate genes that were not detected or were excluded from scoring in the spatial transcriptomics dataset.

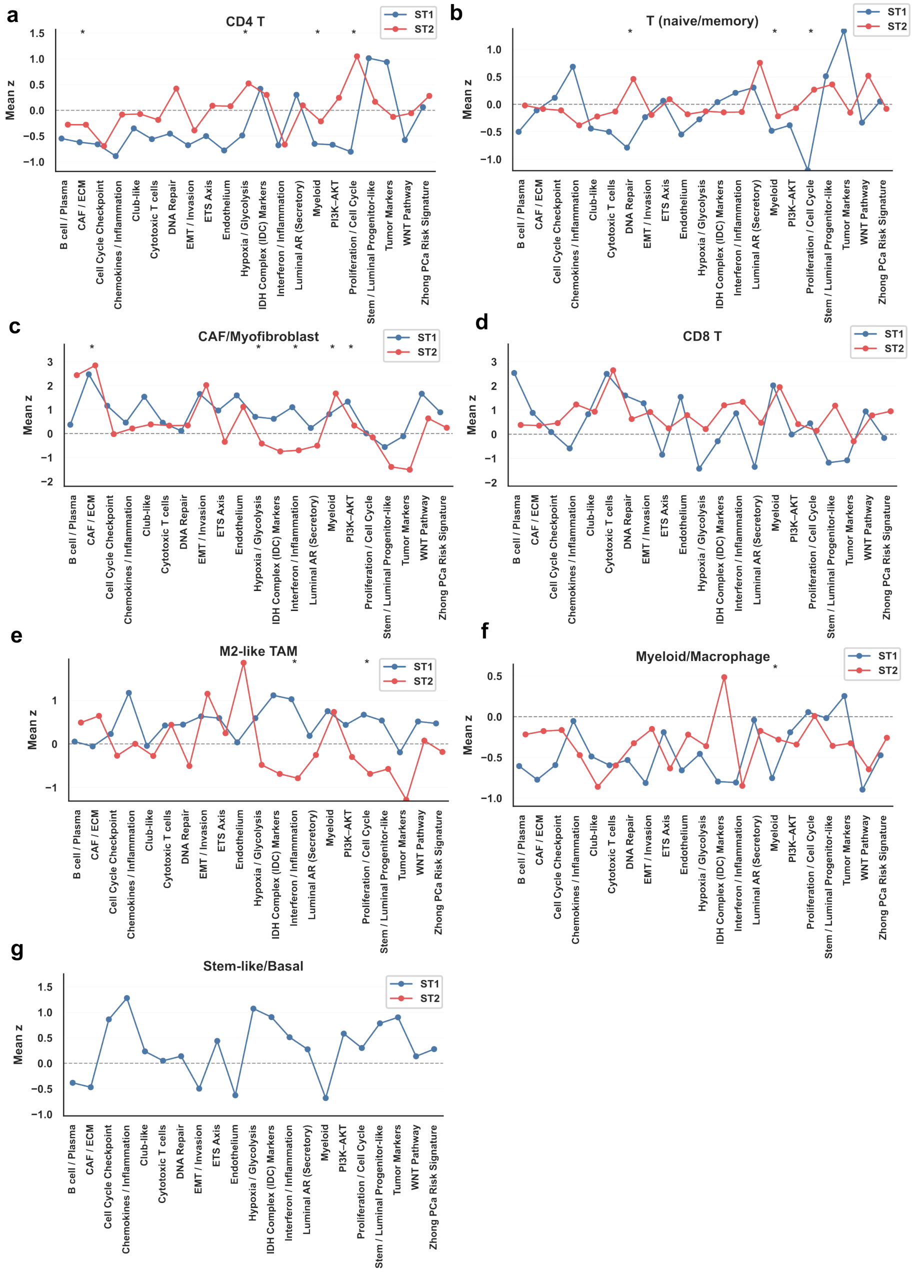


**Supplementary Figure S7: Program-level transcriptional activity across additional cell types in prostate tumour samples.** Program activity profiles comparing prostate sections ST1 (blue) and ST2 (red) for cell types not shown in the main figure. Each point represents the mean z-scored activity of a gene program within the indicated cell type. Horizontal dashed lines indicate the baseline mean program activity, and asterisks denote statistically significant differences based on bootstrap testing. **(a)** CD4 T cells. **(b)** Naive/memory T cells. **(c)** CAF/myofibroblast populations. **(d)** CD8 T cells. **(e)** M2-like tumour-associated macrophages (TAM). **(f)** Myeloid/macrophage populations. **(g)** Stem-like/basal epithelial cells.

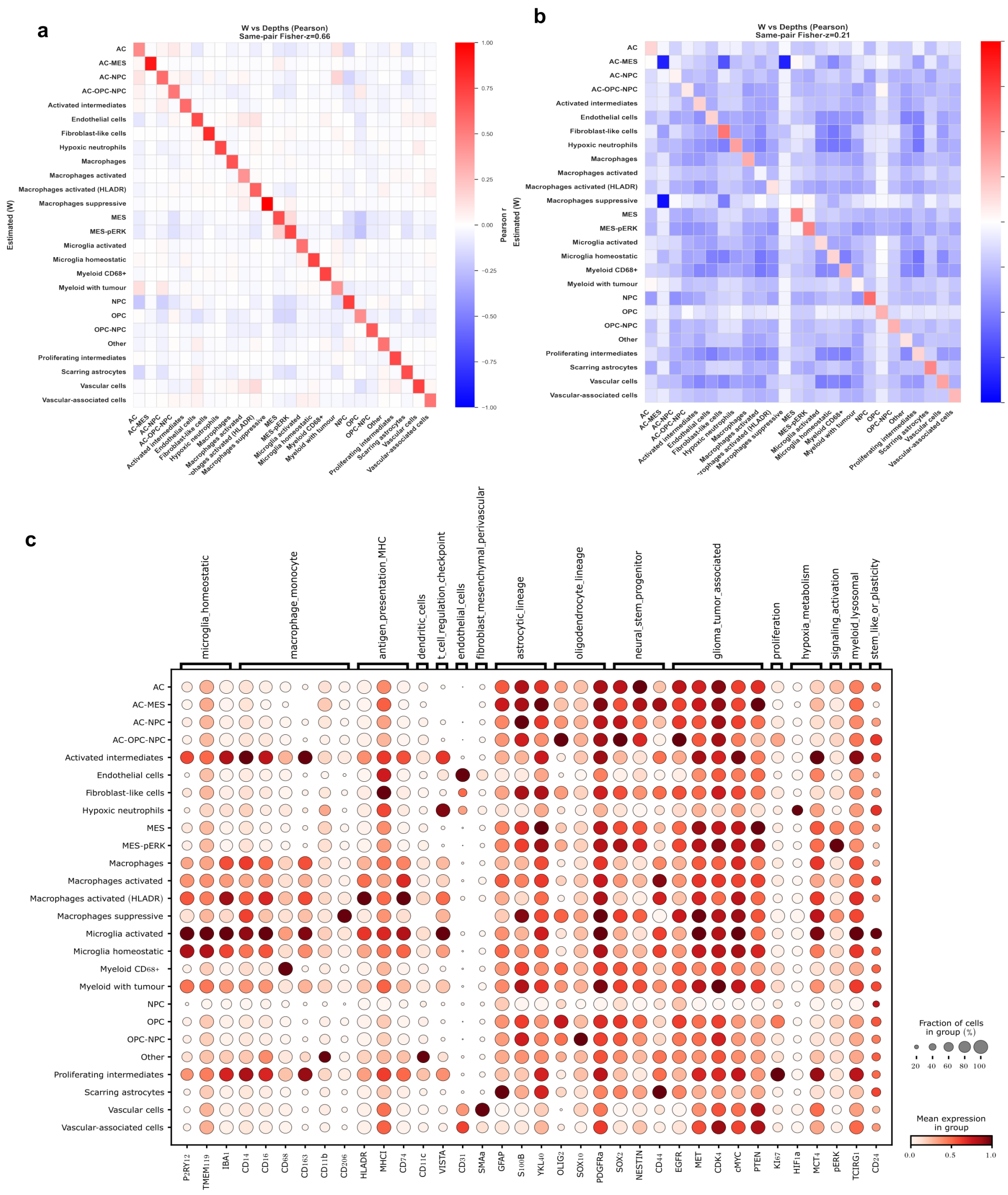


**Supplementary Figure S8: Fine-grained evaluation of inferred GBM cell-type abundances and IMC marker profiles.** **(a)** Pearson correlation between PISTACHIO-inferred cell-type abundances ($W$) and IMC-derived fine-grained ground-truth cell-type depths across spatial spots, computed including shared zero–zero entries. The heatmap shows all pairwise correlations between inferred and ground-truth cell types, with diagonal enrichment indicating correct recovery of individual populations. **(b)** Corresponding correlation analysis excluding shared zero–zero entries, providing a more stringent evaluation focused on spatially informative regions where both inferred and ground-truth signals are present. **(c)** IMC protein marker profiles across fine-grained GBM cell types used to define spatial priors. Dot size indicates the fraction of cells expressing each marker within a cell type, and colour indicates mean protein expression.

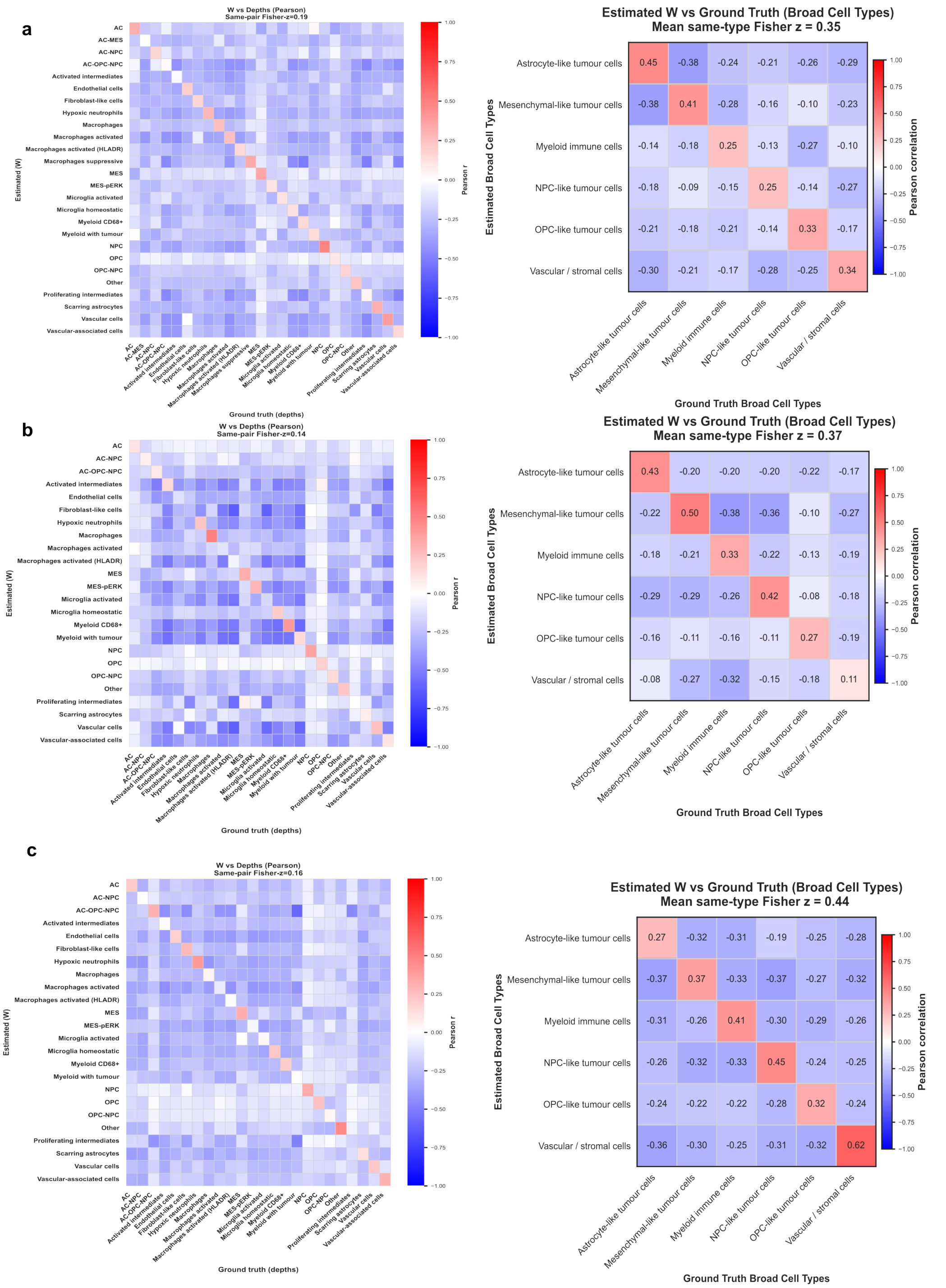


**Supplementary Figure S9: Fine-grained and broad cell-type correlation analyses across GBM samples A–C. (a)** Sample A: Pearson correlation matrices between inferred cell-type abundances ($W$) and IMC-derived ground truth. Left panels show fine-grained cell-type correlations, while right panels summarise correlations after aggregation into broad biological compartments. **(b)** Sample B: Corresponding correlation analyses as in (a), demonstrating recovery of major tumour, immune, and stromal compartments with varying resolution across cell types. **(c)** Sample C: Corresponding correlation analyses as in (a), showing strong diagonal structure at the broad cell-type level and improved separation of major compartments compared with fine-grained annotations. Colour scales indicate Pearson correlation coefficients.

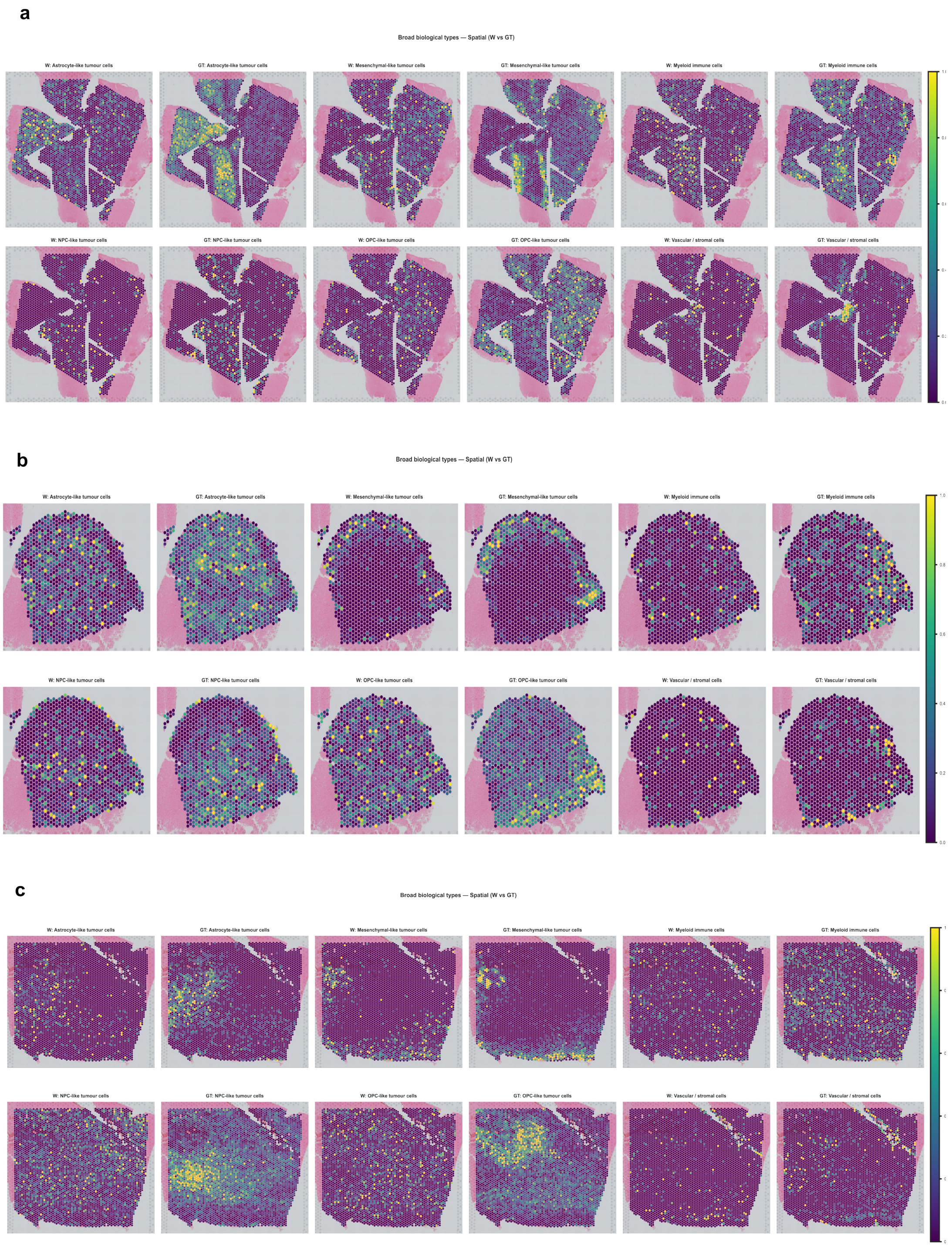


**Supplementary Figure S10: Spatial comparison of inferred and IMC-derived broad cell-type composition across GBM samples A–C. (a)** Sample A: Spatial maps of broad cell-type composition. Columns show IMC-derived ground truth and corresponding spatial maps inferred by PISTACHIO. Rows correspond to astrocyte-like tumour cells, mesenchymal-like tumour cells, myeloid immune cells, NPC-like tumour cells, OPC-like tumour cells, and vascular/stromal cells. **(b)** Sample B: Corresponding spatial comparison as in (a), illustrating recovery of major tumour ecosystems and immune/stromal compartments. **(c)** Sample C: Corresponding spatial comparison as in (a), showing strong agreement between inferred and IMC-derived spatial organisation, particularly for dominant tumour and vascular/stromal regions. Colour scales indicate per-spot cell-type composition using a shared scale within each panel.

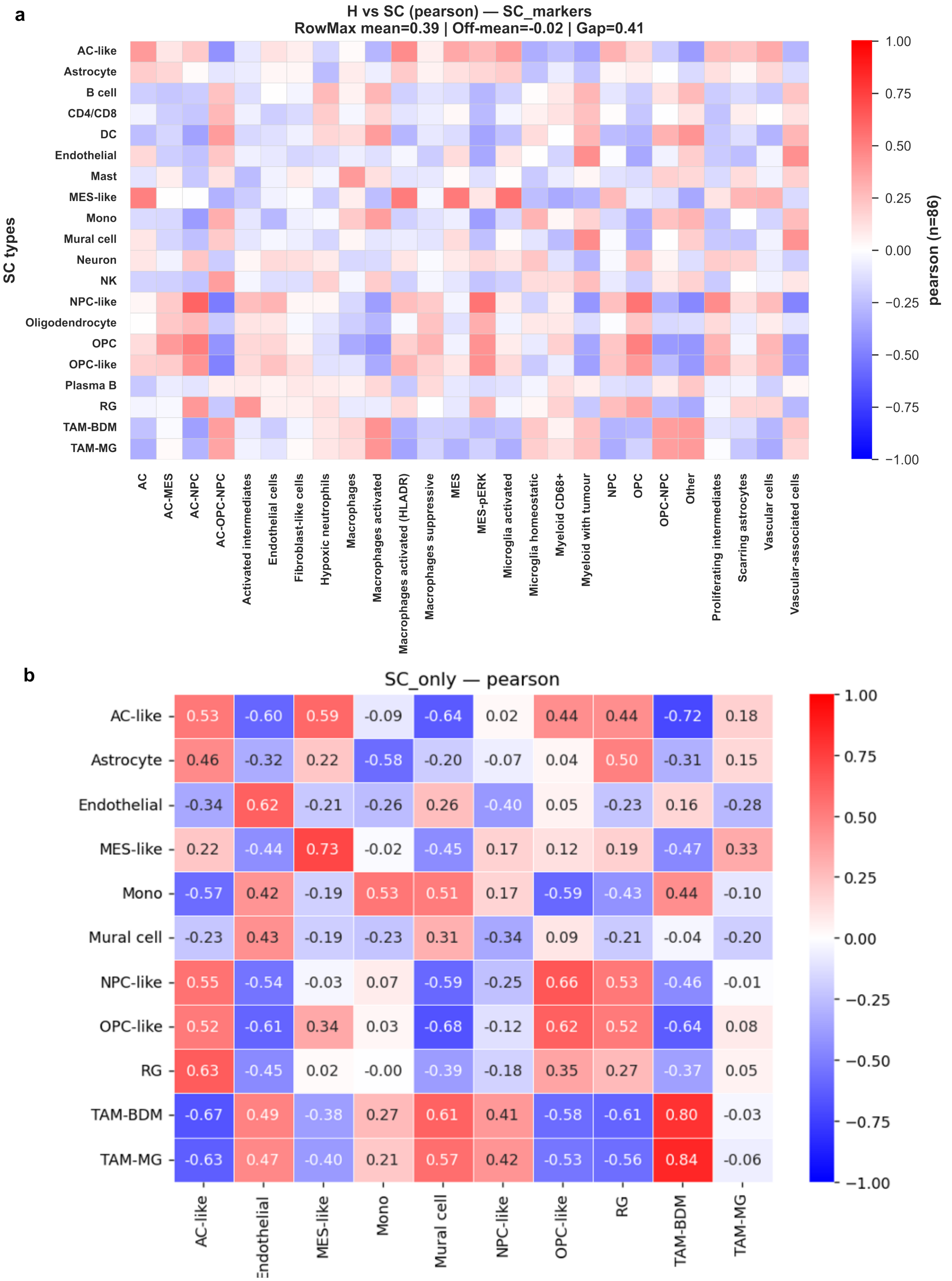


**Supplementary Figure S11: Comparison of inferred GBM transcriptional programs with external single-cell RNA-seq references.** **(a)** Pearson correlation between inferred gene-loading profiles ($H$) and external GBM single-cell reference cell types using curated single-cell marker genes. The heatmap illustrates partial but incomplete correspondence between inferred spatial transcriptional programs and reference RNA-defined cell states. **(b)** Re-evaluation of inferred transcriptional programs after restricting the comparison to RNA cell classes supported by MaxFuse-based protein–RNA correspondences. This matched-cell-type analysis improved the apparent agreement for canonical malignant and myeloid lineages, including MES-like, NPC-like, OPC-like and TAM-related states.

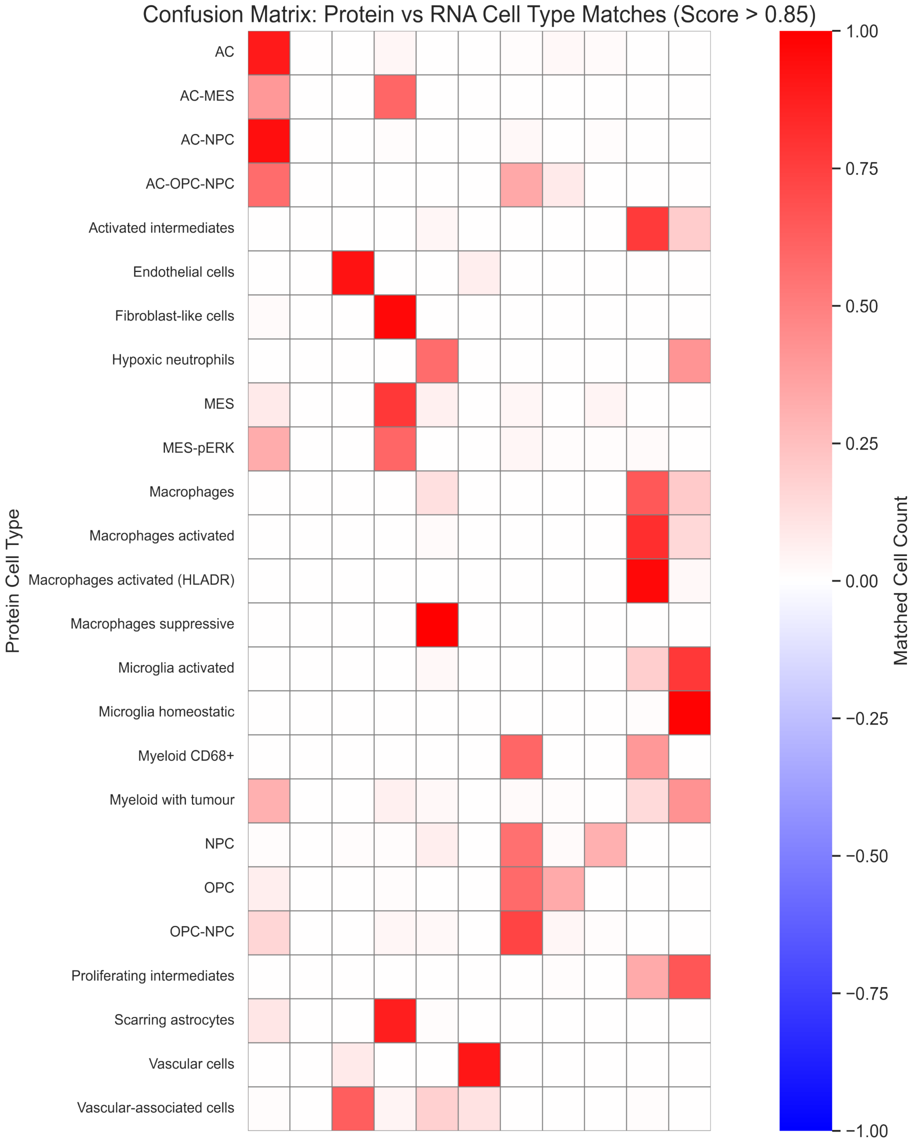


**Supplementary Figure S12: MaxFuse-based correspondence between protein-defined and RNA-defined GBM cell states.** Confusion matrix summarising high-confidence matches between IMC-derived protein cell types and external RNA-defined GBM cell classes. Strong correspondences were observed for canonical astrocyte-like, mesenchymal-like, NPC-like, OPC-like, endothelial, mural and TAM-associated populations, whereas hybrid and transitional protein-defined states mapped across multiple related RNA lineages. These results support partial but incomplete equivalence between protein-based and transcriptome-based annotation systems in GBM.

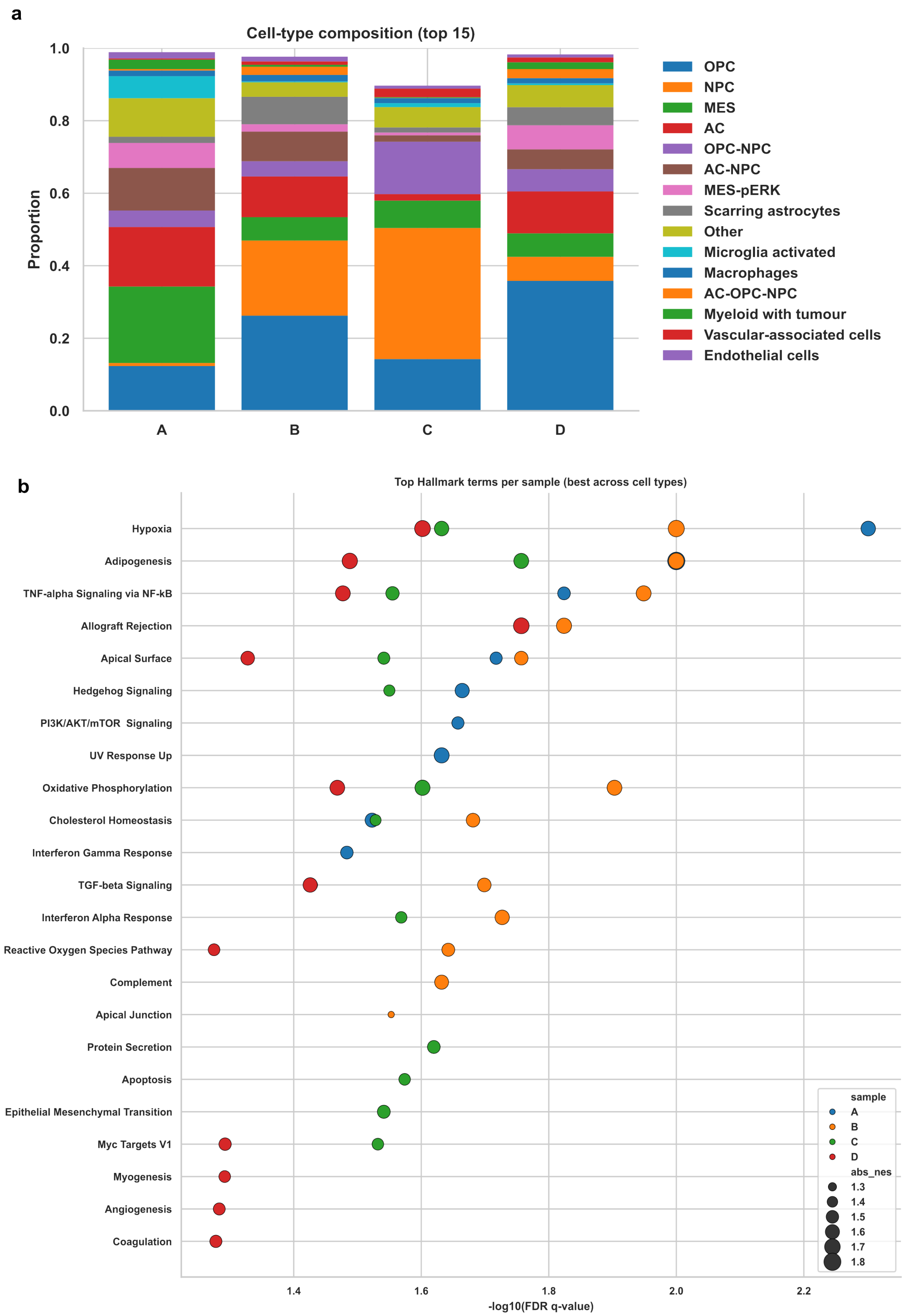


**Supplementary Figure S13: Functional characterisation of inferred GBM transcriptional programs.** **(a)** Broad cell-type composition across GBM samples A–D, shown as proportions of the dominant inferred cell states. **(b)** Top Hallmark terms per sample identified from inferred transcriptional programs, summarised across cell types. Point position indicates enrichment significance ($-\log_{10}$ FDR-adjusted $q$-value), and point size indicates the absolute normalised enrichment score.

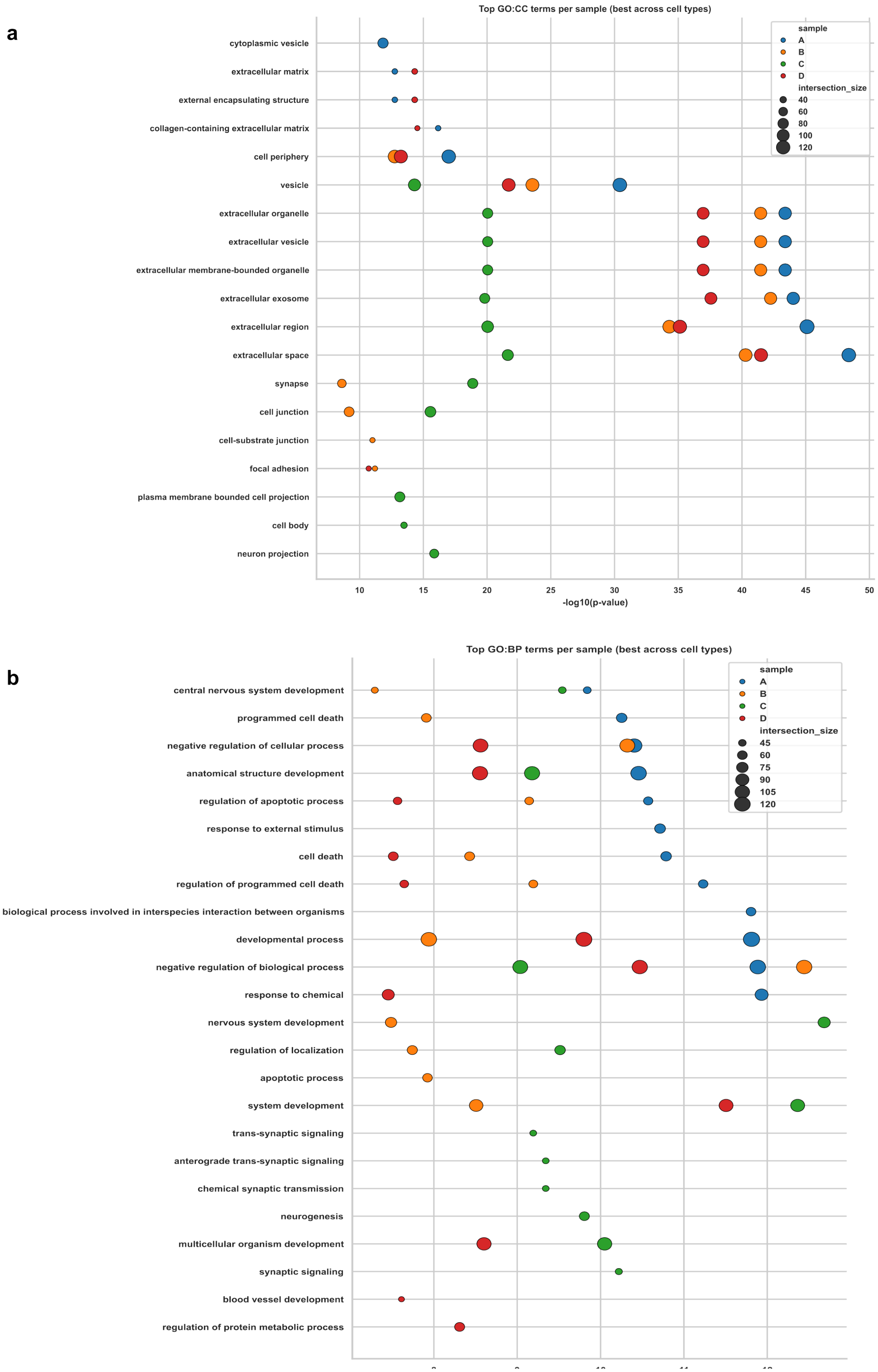


**Supplementary Figure S14: Gene Ontology enrichment of inferred GBM transcriptional programs.** **(a)** Top GO cellular component (GO:CC) terms per sample, summarised across cell types. Enriched terms include extracellular vesicle, extracellular exosome, extracellular region, extracellular matrix, synapse, neuron projection and cell junction-related components. **(b)** Top GO biological process (GO:BP) terms per sample, highlighting nervous system development, neurogenesis, synaptic signalling, developmental processes, apoptotic regulation, response to external stimuli and blood vessel development. Point position indicates enrichment significance ($-\log_{10}$ $p$-value), and point size represents gene-set overlap.

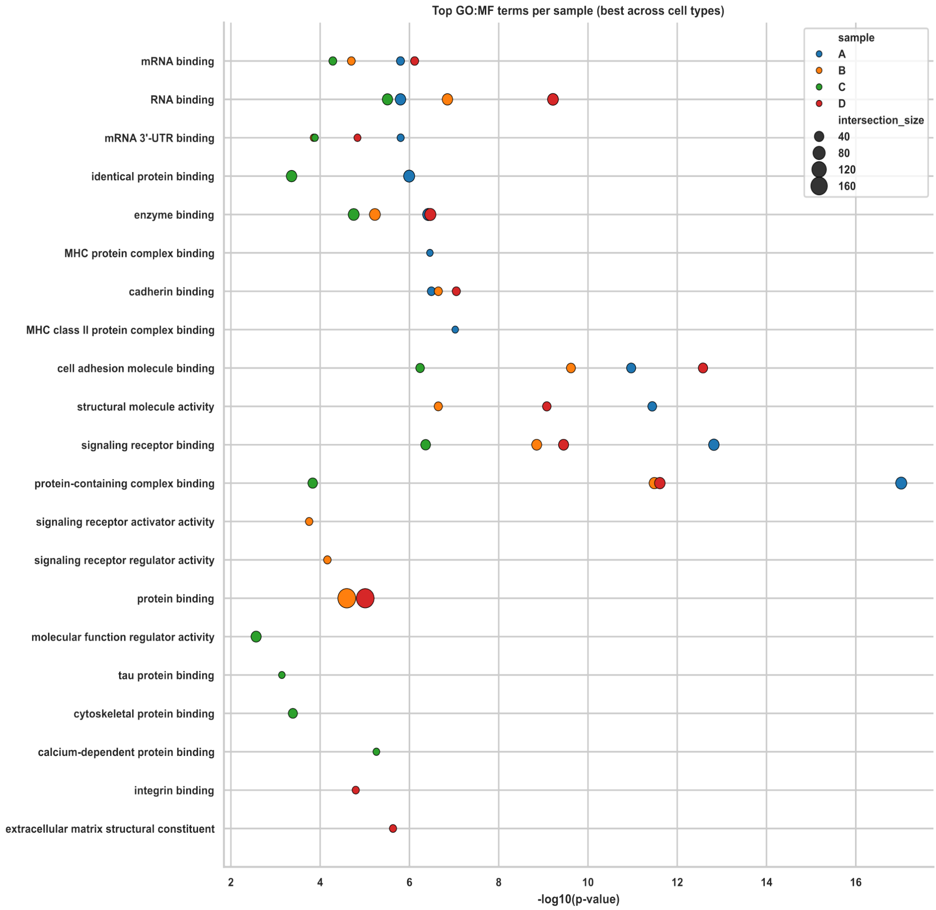


**Supplementary Figure S15: GO molecular function enrichment of inferred GBM transcriptional programs.** Top GO molecular function (GO:MF) terms per sample, summarised across cell types. Enriched functions include protein binding, RNA binding, mRNA binding, cell adhesion molecule binding, signalling receptor binding, cadherin binding, integrin binding and extracellular matrix structural constituent activity, indicating that inferred GBM programs capture signalling, adhesion, matrix interaction and post-transcriptional regulatory processes. Point position indicates enrichment significance ($-\log_{10}$ $p$-value), and point size represents gene-set overlap.
